# Toward Automatic Variant Interpretation: Discordant Genetic Interpretation Across Variant Annotations for ClinVar Pathogenic Variants

**DOI:** 10.1101/2024.10.11.617756

**Authors:** Yu-An Chen, Tzu-Hang Yuan, Jia-Hsing Huang, Yu-Bin Wang, Tzu-Mao Hung, Chien-Yu Chen, Pei-Lung Chen, Jacob Shujui Hsu

## Abstract

**Purpose:** High-throughput sequencing has revolutionized genetic disorder diagnosis, but variant pathogenicity interpretation is still challenging. Even though the Human Genome Variation Society (HGVS) provides recommendations for variant nomenclature, discrepancies in annotation remain a significant hurdle.

**Methods:** This study evaluated the annotation concordance between three tools— ANNOVAR, SnpEff, and Variant Effect Predictor (VEP)—using 164,549 two-star variants from ClinVar. The analysis used HGVS nomenclature string-match comparisons to assess annotation consistency from each tool, corresponding coding impacts, and associated ACMG criteria inferred from the annotations.

**Results:** The analysis revealed variable concordance rates, with 58.52% agreement for HGVSc, 84.04% for HGVSp, and 85.58% for the coding impact. SnpEff showed the highest match for HGVSc (0.988), while VEP bettered for HGVSp (0.977). The substantial discrepancies were noted in the Loss-of-Function (LoF) category. Incorrect PVS1 interpretations affected the final pathogenicity and downgraded PLP variants (ANNOVAR 55.9%, SnpEff 66.5%, VEP 67.3%), risking false negatives of clinically relevant variants in reports.

**Conclusions:** These findings highlight the critical challenges in accurately interpreting variant pathogenicity due to discrepancies in annotations. To enhance the reliability of genetic variant interpretation in clinical practice, standardizing transcript sets and systematically cross-validating results across multiple annotation tools is essential.

**Graphic abstract:** This study examined the consistency of variant annotations produced by three widely used open-source tools—ANNOVAR, SnpEff, and VEP—against 164,549 ClinVar two starts variants. The investigation covers HGVS-based transcript, protein nomenclature and coding impact annotation. The results showed that none of the tools were fully consistent with ClinVar across all coding impact categories, particularly in the LoF category, which exhibited the poorest consistency. This inconsistency may lead to discrepancies in PVS1 interpretation, affecting the final pathogenicity assessment. PVS1 loss resulted in a significant downgrading of PLP variants, potentially leading to the omission of clinically relevant variants in reports.

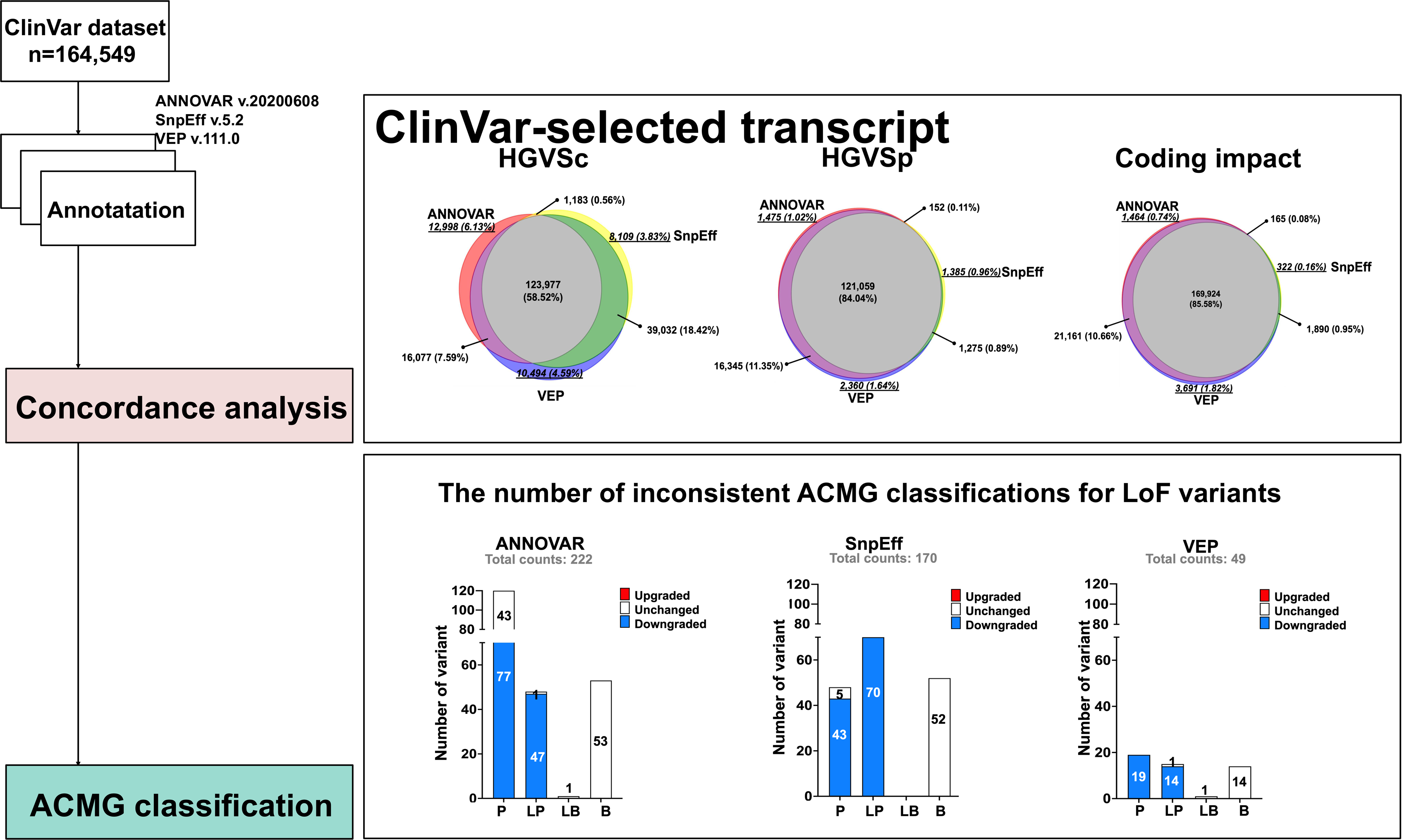

## Background

The application of high-throughput sequencing technology in molecular genetic testing has dramatically improved the research and diagnosis of genetic disorders in clinical practice^1,2^. HGVS provides standards and guidelines for documenting variants at the genomic, transcript (coding), and protein levels^3,4^. With the updated standards and guidelines from the American College of Medical Genetics and Genomics (ACMG) and the Association for Molecular Pathology (AMP) for the clinical interpretation of sequencing variants associated with human diseases^5^, the clinical significance of any given sequence variant could be classified accurately. However, there are still challenges in interpreting variants when translating high-throughput genetic sequencing into clinical practice.

Variant annotation is crucial for linking variants to phenotypic abnormalities and impacts downstream interpretation. Key annotation components include accurately locating variants and assessing their influence on gene products. This involves assigning functional classifications to DNA variants, incorporating sequence conservation metrics^6^, and predicting variant effects on protein structure and function^7,8,9^.

Several databases, such as Ensembl^10^, RefSeq^11^, UCSC^12^, and ENCODE^13^ provide functional characteristics of genomic regions. Annotation tools like ANNOVAR^14^, SnpEff^15^, and VEP^16^ utilize RNA transcripts to annotate variants by using overlapping genomic characteristics. However, coordinating variant positions with transcripts is challenging due to sequence complexities and varying inferred exon structures depending on the bioinformatic tools used.

The MANE project (Matched Annotation from the NCBI and EMBL-EBI) aims to standardize human gene and transcript annotation. MANE transcripts are meticulously selected to precisely match exonic regions between a RefSeq and its Ensembl/GENCODE counterpart^17^. However, different MANE selections still impact the final annotation. For example, an exonic variant on the MANE plus clinical transcript NM_033056.4:c.4699_4715dup may be considered intronic on the MANE select transcript NM_001384140.1 (Figure 1A). Besides, the Variant Call Format (VCF) generated by different callers may result in inconsistent HGVS notation. In cases of nucleotide repeats, shifts to the left or right concerning the gene or transcript can lead to distinct genome positions. For instance, the alteration position of NC_000013.11:g.32316495_32316496del on NM_000059.4 is 31_32 when left-shifted but 35_36 when right-shifted (Figure 1B). Additionally, HGVSp (HGVS protein sequence name) can be expressed with either One-Letter or Three-Letter codes for Amino Acids (aa). The annotations for a variant depend on their alignment, transcript set, and the preference of representing HGVS syntax (Figure 1C). Furthermore, HGVS nomenclature has preferred and non-preferred syntax^4^. For example, the preferred HGVSc (HGVS coding sequence name) of NM_001009944.3:c.5824dup may also be expressed as NM_001009944.3:c.5824_5825ins, as it prefers duplication over insertion. The HGVSp can be either long or short form as NP_001009944.3:p.Arg1942ProfsTer48 or NP_001009944.3:p.Arg1942fs, both are preferred syntax (Figure 1D).

**Figure 1.**
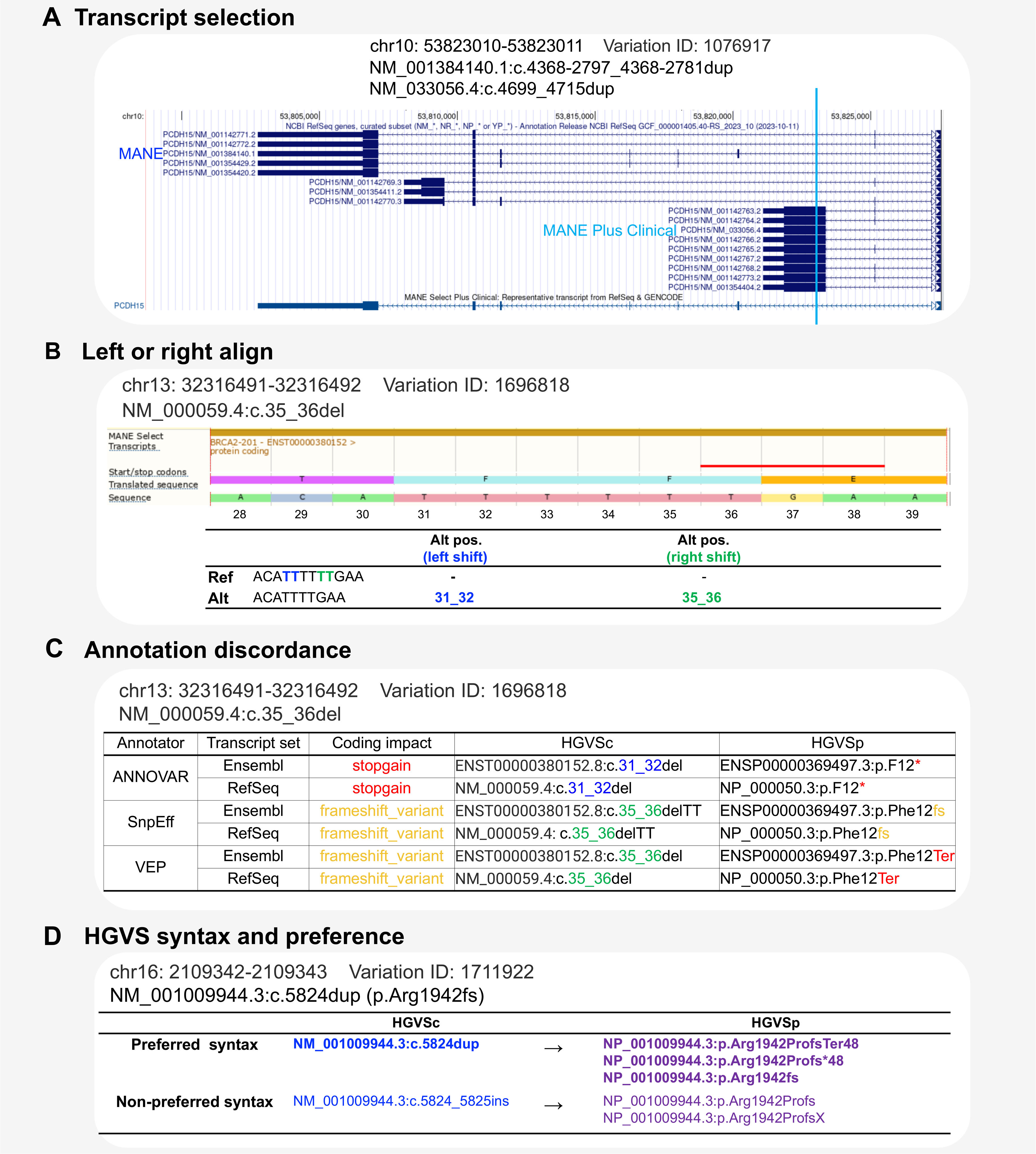
Factors Influencing HGVS Syntax Generation and Coding Impact Annotation. **(A)** The selection of transcript accession affects HGVS syntax, as each accession corresponds to a unique RNA sequence. **(B)** The presence of repeat sequences allows for two-directional alignment (right or left), which can impact the final HGVS expression and associated annotation. **(C)** Annotation outputs from (B) are presented, highlighting discrepancies in the results. **(D)** An illustration showing that HGVS syntax for the same coding variant can have multiple synonyms, with HGVS specifying preferred syntax for identical variants.

Previous comparisons noted the significant annotation differences between ANNOVAR and VEP in the chosen transcripts^18^. However, comprehensive evaluations of consistency with reputable clinical databases are lacking, and the impact on ACMG interpretation needs to be explored. With the updated annotation tools, we aimed to reassess the variant interpretation accessibility, comparing the concordance of variant nomenclature and predicted coding impacts generated by ANNOVAR, SnpEff, and VEP for 164,549 high review status ClinVar variants^19,20^. We evaluated the clinical implications according to ACMG guidelines and proposed critical points to improving the implementation of genetic sequencing into routine clinical practice.

## Methods

### Datasets

The VCF file (version January 7, 2024) was obtained from ClinVar^21^ and underwent preprocessing using bcftools^22^. This preprocessing involved left alignment, removing duplicates, eliminating degenerate bases, filtering out variants with no alternative alleles, and normalization. Subsequently, the dataset underwent thorough curation following stringent filtering criteria (Figure 2A). A total of 164,549 variants were selected from the ClinVar VCF^23^.

**Figure 2.**
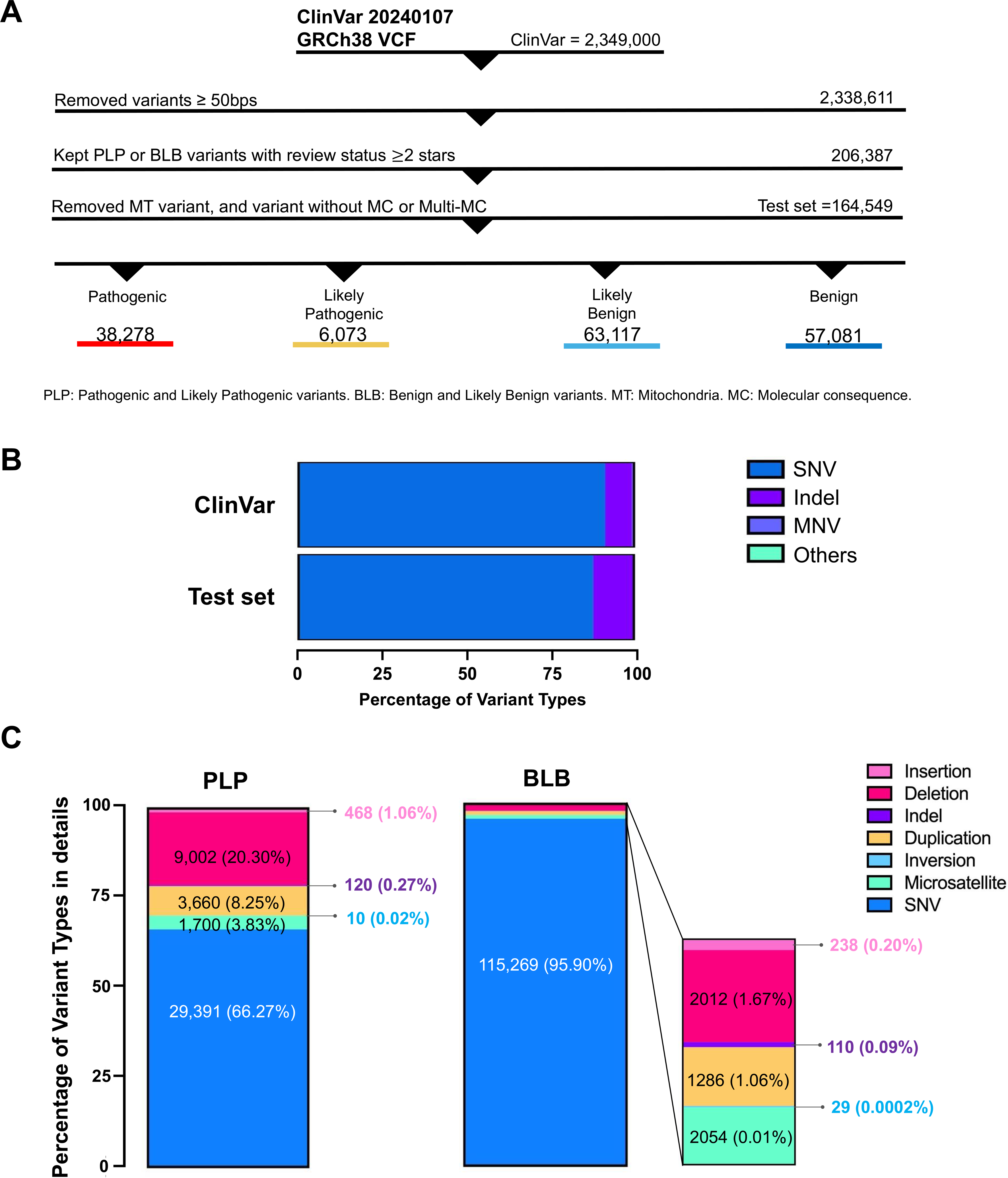
Input Dataset Characteristics and Composition. **(A)** Illustration of the dataset preprocessing and the ClinVar pathogenicity classification, including PLP (Pathogenic and Likely Pathogenic variants), BLB (Benign and Likely Benign variants), MT (Mitochondria), and MC (Molecular Consequence). **(B)** The stacked bar showing the number of variants and distribution of variant types, with the x-axis representing the Percentage of Total Variants. SNV: single nucleotide substitution. Indel: insertion/deletion. MNV: multi-nucleotide variant. **(C)** The stacked bars depicting the composition of variant types in the PLP and BLB variant subsets, with the y-axis representing the Percentage of Variant Composition. Due to discrepancies in transcript annotations, the number of evaluated variants may be less than the total number of variants in the input set.

### Variant annotations & Sequence Ontology (SO) normalization

Annotations for HGVS and coding impact utilized by ANNOVAR^14^ (version June 8, 2020), SnpEff^15^(version 5.2, released April 9, 2024), and VEP^16^ (version111.0). All tools supported VCF input; the genome build was NCBI GRCh38p13. Variant annotations were obtained using integrated RefSeq and ENSEMBL transcript sets, detailed in Key Resource Table “Software and dataset used in this study.”^14,15,16,22,23,24,25,26,36^

In ANNOVAR, we adjusted the upstream and downstream parameters from the default 1000 bp to 5000 bp to ensure consistency with the other two tools, and we enabled HGVS output. For VEP, we enabled HGVSg, HGVSc, and HGVSp. SnpEff was used with default settings. The outputs were normalized to standardize the terms and prioritize the most severe impact for our evaluation (Supplementary Table S1). A comparison table was produced with an R script from VCF files containing ANNOVAR, SnpEff, and VEP annotations and gene information for the transcript(s) used for each annotation.

### Syntax Comparisons of Variant Annotations

We performed string-match comparisons between the output and the reference syntax with variants described on the exact transcript accession (Additional file 2: Figure S1). Variant annotations were considered matches when the HGVS string and the query annotation matched as-is. The tool’s annotation was also considered a match if the string did not match perfectly but could be an equivalent expression.

### Automatic ACMG classification

Each inconsistent variant (between each tool and ClinVar) was annotated using an in-house developed workflow Gendiseak (GDK) platform^26^ based on ACMG rules from the 2015 guideline^5^, with adjustments to specific rules based on internal testing. Specifically, for PP5 and BP6, variants with a ClinVar review status of two-star were adjusted to PP5-moderate and BP6-strong, and those with ≥ three-star to very strong. PP3 was disabled when PVS1 was triggered. For PP3, we followed ClinGen recommendations^27^, with thresholds set at CADD>25.3 or SpliceAI>0.5. Rules related to gene-disease relations or requiring cohort study or phasing information (PS4, PM3, PP4, BP2, BP5) were excluded.

Firstly, we interpreted ACMG classes based on ClinVar’s LoF coding impact. To simulate an automated interpretation scenario, the variants with coding impact belonging to the LoF category were assigned PVS1 directly. Variant classifications based on ClinVar’s LoF coding impact and the other rules without conflicts with ClinVar records were classified as “no conflict.” Subsequently, these “no conflict” variants were interpreted based on the coding impact of each annotator. We assessed the number of LoF variants for which discrepancies between annotators’ and ClinVar’s coding impact led to ACMG classification changes.

### Plotting and statistical analysis

Venn diagrams in this manuscript were generated by Python (version 3.9) with matplotlib-venn^28^ GraphPad Prism™ software^29^ (version 7.0a) was used to analyze results and plot the bar charts.

## Results

### 164,549 curated ClinVar Variants

The VCF file (version January 7, 2024) was obtained from ClinVar^21^ and curated following stringent filtering criteria. Of the 2,349,000 variants in the ClinVar VCF^23^, most are SNVs (2,147,692; 91.43%), with fewer indels (186,883; 7.96%), MNVs (7,847; 0.33%), and other types (6,578; 0.28%) (Figure 2A, 2B). Specifically, we excluded indels ≥ 50 bp to analyze small variants. To ensure reliable pathogenicity information, we retained variants with a ClinVar review status of ≥ two-star and excluded variants classified as variants of uncertain significance (VUS). We also excluded mitochondria (MT) variants, those lacking molecular consequence (MC) information, and multi-MC for a single transcript to facilitate subsequent analysis based on ACMG guidelines. Finally, we have a test set with 164,549 variants, composed of SNVs 144,660 (87.91%), 19,620 indels (11.92%), and 269 MNVs (multi-nucleotide variants) (0.16%). The composition of test set is composed of 38,278 pathogenic (P) variants, 6,073 likely pathogenic (LP) variants, 57,081 benign (B) variants, and 63,117 likely benign (LB) variants. The composition of PLP and BLB was shown separately, and the variant type of PLP is more diverse than that of BLB (Figure 2C), implying that the impact of annotation may be more significant in PLP variants.

### Transcript Availability in ANNOVAR, SnpEff, and VEP

The test set was subsequently analyzed using the latest versions of ANNOVAR (version June 8, 2020), SnpEff (v5.2), and VEP (v111.0). Variant annotation was conducted using both RefSeq and Ensembl transcript sets based on GRCh38p13. In this study, we only consider transcripts with annotations available under the categories NM (RefSeq mRNA), NR (RefSeq non-coding RNA), and ENST (Ensembl Transcript). Since a single transcript may harbor multiple variants, and a single variant can correspond to multiple transcripts, the transcripts included in this count are those for which each annotator can provide annotations. Initially, we collected all available transcripts from each tool. VEP had the highest available transcripts (RefSeq: 25,381; Ensembl: 67,695), followed by ANNOVAR (RefSeq: 20,285; Ensembl: 27,148), with SnpEff (RefSeq: 5,848; Ensembl: 6,183) having the fewest. SnpEff’s lower number of available transcripts was partly due to its most recent version, which only included updated MANE selection transcripts. Despite identical input variants, the tools generated different numbers of transcripts and annotations (Figure 3A, B).

**Figure 3.**
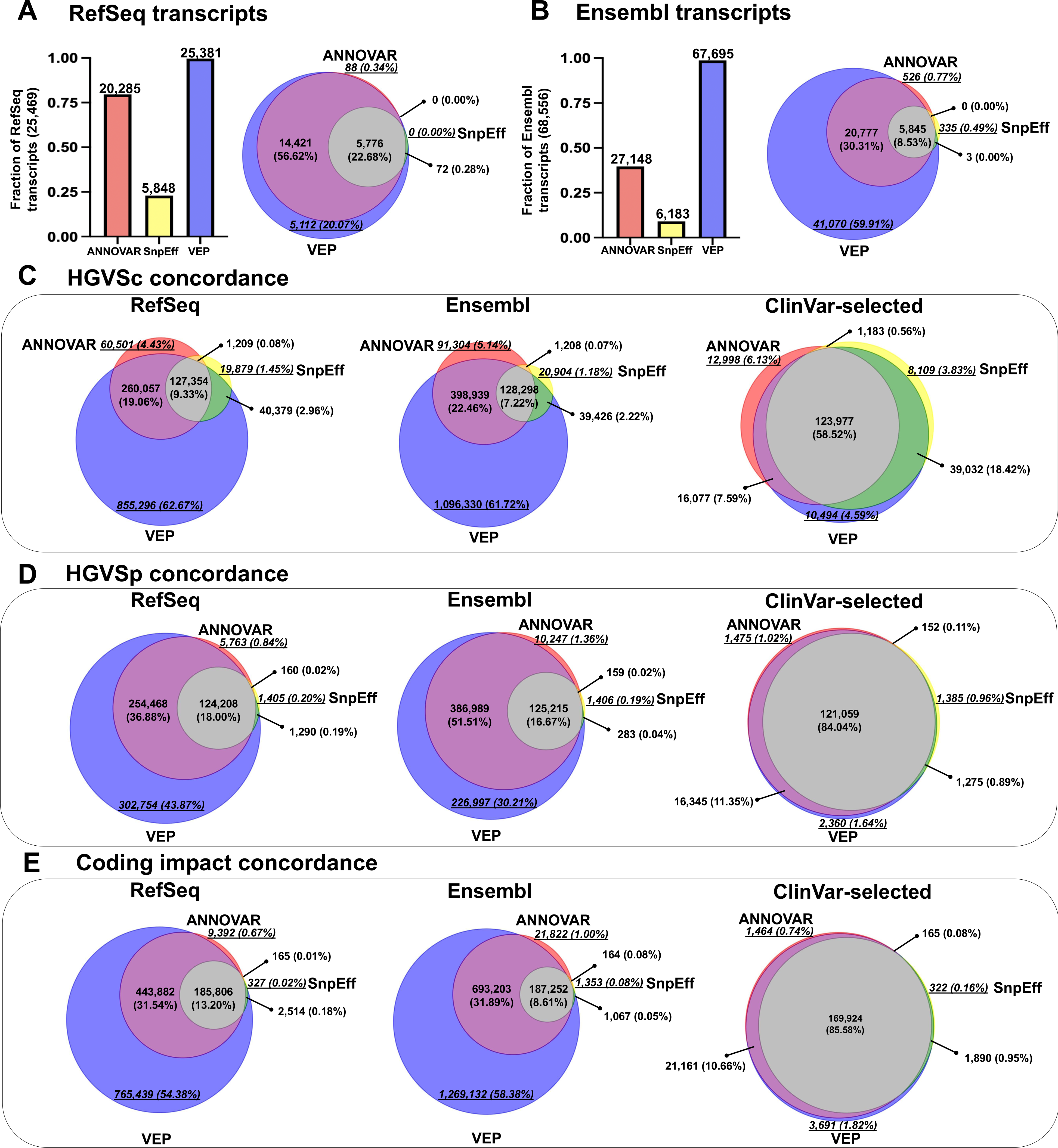
Concordance of HGVSc, HGVSp, and Coding Impact. **(A)** Fraction of each transcript set in relation to the union of all RefSeq transcripts. **(B)** Fraction of each transcript set compared to the union of all Ensembl transcripts. **(C)** Venn diagram illustrating the concordance of HGVSc syntax among ClinVar test-set variants (N=164,549) annotated by ANNOVAR, SnpEff, and VEP, against the ClinVar records. The comparisons were made based on RefSeq, Ensembl and ClinVar-selected transcript sets. **(D)** Venn diagram depicting the concordance of HGVSp syntax. **(E)** Venn diagram showing the concordance of protein coding impact consequences. The Venn diagram illustrates the overlaps (Red: ANNOVAR only, Yellow: SnpEff only, Purple: VEP only, Orange: ANNOVAR ∩ SnpEff, Green: SnpEff ∩ VEP, Magenta: ANNOVAR ∩ VEP, and Gray: ANNOVAR ∩ SnpEff ∩ VEP). Tool-specific numbers and proportions are denoted in italics and underlined.

Transcript concordance was evaluated by matching transcript accessions. The concordance of RefSeq and Ensembl transcripts among the three tools was 22.68% (n=5,776) and 8.53% (n=5,845), respectively. VEP had the highest number of unique transcripts (RefSeq: 5,112; Ensembl: 41,070) compared to ANNOVAR (RefSeq: 88; Ensembl: 526) and SnpEff (RefSeq: 0; Ensembl: 335) (Figure 3A, B). Inconsistent transcript numbers suggest that annotating the variant’s impact may only be comparable if a transcript is available, highlighting the importance of transcript set selection.

### 58.5% Concordance in HGVSc Syntax for ClinVar-Selected Variants

Defining a variant’s location on a transcript and its potential effect is fundamental to interpretation. Researchers rely on HGVS as a string to search databases or literature, considering ACMG rules such as PS1/PM5, PS3/BS3, PP5/BP6. The coding impact predictions also aid in interpreting specific categories of variants, such as LoF variants for PVS1, missense for PM1 and PP2/BP1, protein length-change for PM4/BP3, and synonymous variants for BP7. Given the importance of HGVS and coding impact annotation in ACMG interpretation, we aimed to evaluate the consistency of HGVS expressions and variant coding impact among annotation tools. The ACMG rules discussed in this study are listed in Additional file 1: Table S2.

Comparing HGVS syntax equivalency from different tools is challenging due to variations in HGVS formatting. Each tool has its unique format, and the alternative forms of HGVS add complexity. For instance, frameshift variants can be represented as p.Phe12LeufsTer13, p.Phe12fs, and F12Lfs*13, while synonymous variants can be described as p.Val17=, p.Val17Val, and p.V17V. The annotated outputs often did not exactly match the preferred HGVS syntax; for instance, ANNOVAR and SnpEff represent synonymous variants such as NP_003997.1:p.Cys188Cys, whereas the HGVS preference is NP_003997.1:p.Cys188=. For nonsense variants, ANNOVAR denotes them as p.Q480X, while the HGVS preference is p.Gln480Ter or p.Gln480*. The exemplar variants demonstrating nomenclature discrepancies of HGVSc and HGVSp are shown in Table S3 and Table S4.

To assess the equivalency of HGVS nomenclature across tools, we focused on HGVSc and HGVSp. We conducted string-match comparisons between the output and the reference syntax for variants described on the exact transcript accession (Additional file 2: Figure S1). The required format is “Position + Accession number of transcripts (NM or ENST) + HGVSc syntax.” For example, intergenic variants, which do not have HGVSc annotations, are not included in the comparison. Since ANNOVAR only provides transcript accession information without specifying versions, we considered a match between the query transcript and the reference transcript accession for HGVSc string-match comparison, ignoring version numbers (Additional file 2: Figure S1A). If the transcript accessions differed, it was considered incorrect. For HGVSp string-match comparison, because ANNOVAR did not provide “NP” or “ENSP” accession numbers, we compared the coding DNA transcript accession and protein symbol. Then, we compared the HGVSc and HGVSp syntax of both the query and reference expressions on the same transcript based on HGVS recommendations (Additional file 2: Figure S1). If the syntax for both expressions was equivalent, it was considered correct. It was considered incorrect if the syntax differed in position, variant type, or was empty.

The concordance of HGVS syntax between tools was analyzed and shown in Figure 3C and 3D. The concordance was low, with only 9.33% and 7.22% agreement on HGVSc for RefSeq and Ensembl transcripts, respectively, and 18.00% and 16.67% for HGVSp. The differences in the number of available transcripts influenced the results, as a missing transcript was considered incorrect, highlighting the importance of transcript usage in annotation tools. In clinical practice, including the desired transcript set and selecting the most clinically relevant transcripts is crucial for the interpretation.

For SnpEff, which only supports MANE transcripts, we specifically compared HGVSc and HGVSp annotations for ClinVar-selected transcripts, most of which are in MANE-selected transcripts. The concordance between tools was higher in ClinVar-selected transcripts, with 58.52% agreement on HGVSc and 84.04% on HGVSp. However, we still observed discordance in HGVS expression across tools. ANNOVAR showed the most differences in HGVSc, with 12,998 (6.13%) unique syntaxes (Figure 3C), while VEP had the most differences in HGVSp, with 2,360 (1.64%) unique syntaxes (Figure 3D).

Although HGVS syntax from different tools can be equivalent in molecular consequence type, variations in syntax can pose challenges when searching literature or databases, especially with automated NGS variant annotations. Inaccurate variant-disease relation information may impact the final ACMG interpretation. These results highlight the inconsistencies in HGVS representation, leading to confusion and uncertainty in variant interpretation, and emphasize the need to carefully consider transcript selection and HGVS syntax accuracy when using annotation tools to ensure precise variant interpretation.

### 85.6% Agreement in Coding Impact for ClinVar-Selected Variants

As variant interpretation based on ACMG guidelines often relies on coding impact information to determine pathogenicity, we compared the concordance of coding impact annotations among the tools. Since each tool may use different naming conventions, we employed Sequence Ontology (SO) terms to standardize the naming conventions of the three annotators (Additional file 1: Table S1). Besides, since annotation tools may provide multiple annotations for a single variant in a given transcript, we opted to prioritize the most severe impact of the variant in the output for our evaluation. For example, ClinVar variant ID:1483650, NC_000001.11:g.45330514del, is noted in the MANE Select transcript as NM_001048174.2:c.1434+2del, classified as a splice donor variant. In the MANE Plus Clinical annotation, it is noted as NM_001128425.2:c.1518+2del, also classified as a splice donor variant. However, SnpEff annotated this variant as both a splice donor variant and an intron variant in both the MANE Select and MANE Plus Clinical transcripts. Therefore, we have chosen the severe impact splice donor as the representative classification for SnpEff. The concordance of coding impact between tools was analyzed and depicted in Figure 3E. The results revealed that the concordance of coding impact annotations between the three tools was only 13.2% and 8.61% consistent based on RefSeq and Ensembl transcripts, respectively. They had a much higher agreement, 85.58%, based on ClinVar-selected transcripts. Our results indicated that even when based on ClinVar-selected transcripts, there remains a proportion of inconsistency among tools in coding impact annotations. These results demonstrate the importance of prioritizing transcripts and coding impact in handling the large volume of interpretations in clinical testing. These inconsistencies may affect subsequent pathogenicity interpretation, especially for the current ACMG guidelines, as the above rules correspond to each category. For example, ClinVar variation ID: 1879044, NM_000419.5:c.310+3_310+6del, with a review status of ‘Reviewed by expert panel’ curated by ClinGen Platelet ACMG Specifications v2-1, is highly specific for Glanzmann thrombasthenia. The coding impact of this variant in ClinVar is splicing variant, but in ANNOVAR, SnpEff, and VEP, it was annotated as an intron variant.

### High Concordance for SNV and Low Concordance for LoF Variants

Next, we assessed consistency in different variant types and the corresponding functional categories for each annotation tool. For the HGVS comparison, only variants with coding or protein transcript and HGVS syntax information from ClinVar were included, following the string-match comparison rule. We performed SO normalization and prioritization for coding impact annotations as detailed in Additional file 1: Table S1. The fraction of HGVSc matching ClinVar is highest with SnpEff (0.988), followed by VEP (0.977), and then ANNOVAR (0.753), performing the worst (Figure 4A). The discrepancy is less pronounced at the protein level. Regarding HGVSp, VEP demonstrated the highest fraction of matching ClinVar (0.991), followed by SnpEff (0.986) and ANNOVAR (0.981) (Figure 4A).

**Figure 4.**
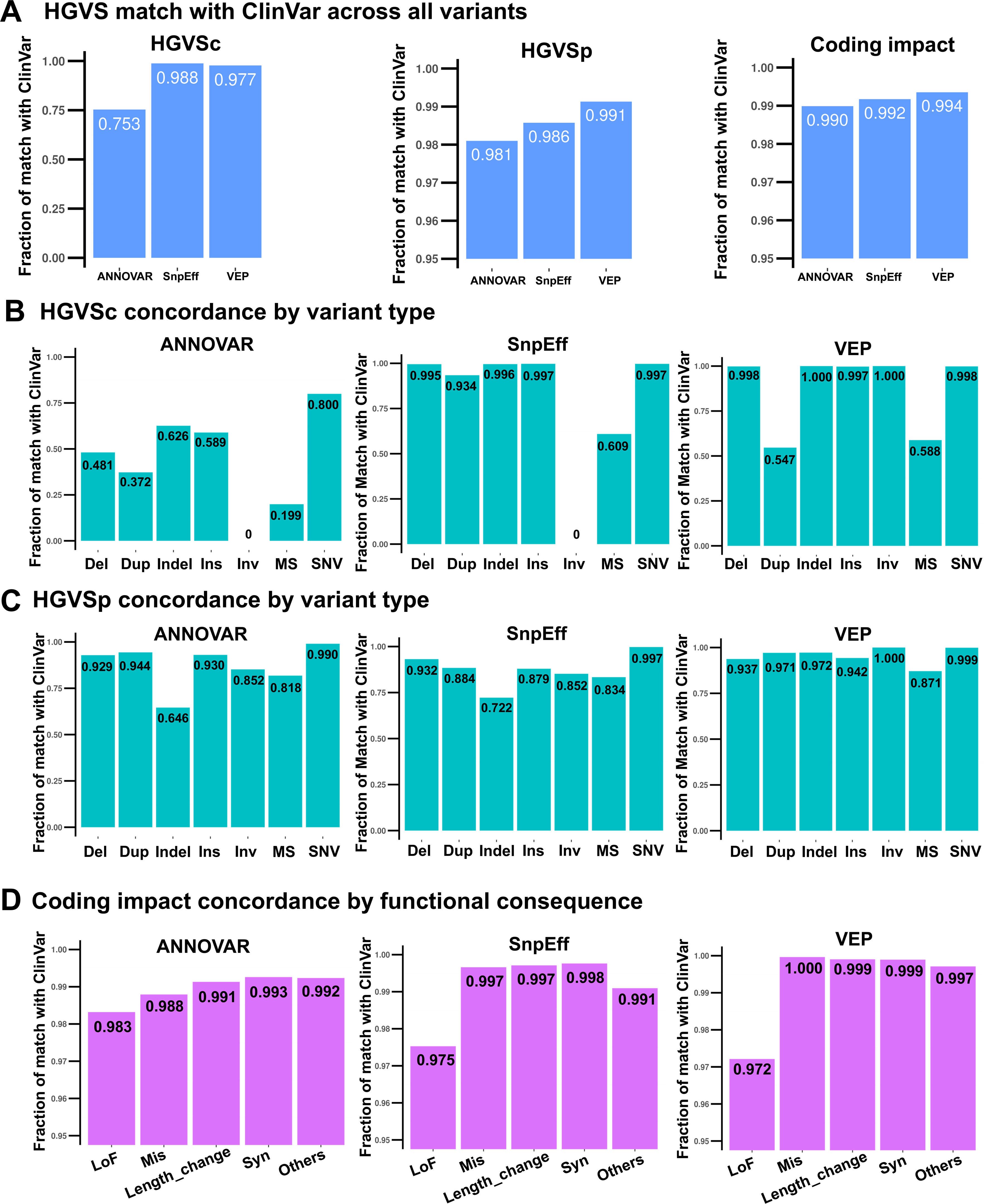
Annotation Match with ClinVar Across Tools. **(A)** Histogram showing the concordance of HGVSc, HGVSp, and Coding impact annotations as percentages of matches with ClinVar records. **(B)** Histograms of HGVSc concordance among the three tools by variant type, highlighting the sensitivity of each tool in capturing HGVSc syntax. **(C)** Histograms of HGVSp concordance among the three tools by variant type. **(D)** Histogram comparing coding impact consequences categorized by functional impact among the three tools. Del: deletion; Dup: duplication; Indel: insertion-deletion; Ins: insertion; Inv: inversion; MS: microsatellite; SNV: single-nucleotide variant; LoF: loss of function; Mis: missense; Syn: synonymous.

We discovered that the majority of inconsistencies between HGVSc annotations from ANNOVAR and ClinVar were due to missing (NA) annotations (79.19%), with the remaining discrepancies attributed to differences in position or variant type (19.67%) or labeled as unknown (1.14%). For SnpEff, HGVSc inconsistencies with ClinVar were primarily due to differences in position or variant type (78.00%) and missing (NA) annotations (22.00%). Similarly, for VEP, HGVSc discrepancies with ClinVar were primarily due to differences in position or variant type (95.47%) and missing (NA) annotations (4.53%) (Additional file 2: supplementary Figure S2). As for HGVSp, the majority of inconsistencies between HGVSp annotations from ANNOVAR and ClinVar were due to differences in position or variant type (57.75%), with the remaining discrepancies attributed to missing (NA) annotations (41.22%) and non-preferred syntax (1.03%). For SnpEff, HGVSp inconsistencies with ClinVar were primarily due to differences in position or variant type (80.11%), followed by missing (NA) annotations (18.51%) and non-preferred syntax (1.37%). For VEP, HGVSp discrepancies with ClinVar were primarily due to HGVS differences in position or variant type (92.89%), followed by missing (NA) annotations (6.55%) and non-preferred syntax (0.56%). (Additional file 2: supplementary Figure S2).

We also assessed the consistency by variant types as follows: Del (deletion), Dup (duplication), Indel (insertion-deletion), Ins (insertion), Inv (inversion), MS (microsatellite), and SNV (single-nucleotide variant). All tools demonstrated the highest consistency with ClinVar for SNV at the coding level compared to other variant types, especially SnpEff and VEP, which yielded near-perfect concordance for SNV. However, the consistency for MS was substantially lower among all variant types in all tools (Figure 4B). Except ANNOVAR, SnpEff, and VEP matched ClinVar with fractions over 0.995 for Ins, Del, and Indel (Figure 4B). SnpEff performed the worst in Inv, while VEP performed the worst in MS and Dup. ANNOVAR showed inferior consistency across almost all categories compared to the other tools.

Similarly, SNVs showed good agreement regarding protein level, but discrepancies were observed in other variant types (Figure 4C). SNVs consistently demonstrate the highest consistency with ClinVar across all tools. Overall, VEP performed best across all variant types, with match fractions to ClinVar of nearly 0.900 or higher for most categories, except for MS (0.871). ANNOVAR and SnpEff also exhibit uniform performance across all variant types, with match fractions to ClinVar of nearly 0.850 or higher for most categories, except for Indel (ANNOVAR: 0.646 and SnpEff: 0.722) and MS (ANNOVAR: 0.818 and SnpEff: 0.834) (Figure 4C). Several rules in the ACMG guidelines require referencing database information; using HGVS syntax search is common. Missing HGVS annotations or inconsistencies in position or variant type may lead to these rules being misinterpreted or incorrectly applied.

The concordance of coding impact annotations was evaluated, showing high agreement between tools and ClinVar in all tools (ANNOVAR: 0.990, SnpEff: 0.992, and VEP: 0.994) (Figure 4A). To explore the coding impact on variant interpretation, we categorized the impact according to ACMG rules into the following categories: 1. Loss of Function (LoF): frameshift, splicing donor, splicing acceptor, and nonsense variants. 2. Missense: missense variants. 3. Protein Length Change: inframe indel, stop-loss, and initiator codon variants. 4. Synonymous: Synonymous variants. 5. Other: variants in UTR, intergenic, upstream/downstream, non-coding (nc), and intronic regions. The detailed mappings of each functional category to the corresponding ACMG rule are listed in Additional file 1: supplementary Table S2. Among variant functional categories, we observed that the LoF category has the lowest agreement (Figure 4D) for all tools (N=31,640, ANNOVAR: 0.983, SnpEff: 0.975, VEP:0.972).

### Impact of Discrepant Coding Annotations on ACMG Classification

All tool-specific discrepancies in functional categories arising from inconsistencies in coding impact annotations have been investigated. Overall, ANNOVAR has the most discrepancies, followed by SnpEff and VEP (Figure 5A, ANNOVAR: 1,666, SnpEff: 1,363, VEP:1,037). Our analysis then focused on whether the coding impact annotation assigned by each annotation tool resulted in a category change. A nonsense variant in ClinVar and frameshift by another tool stays in the LoF category, but a missense annotation shifts it to missense.

**Figure 5.**
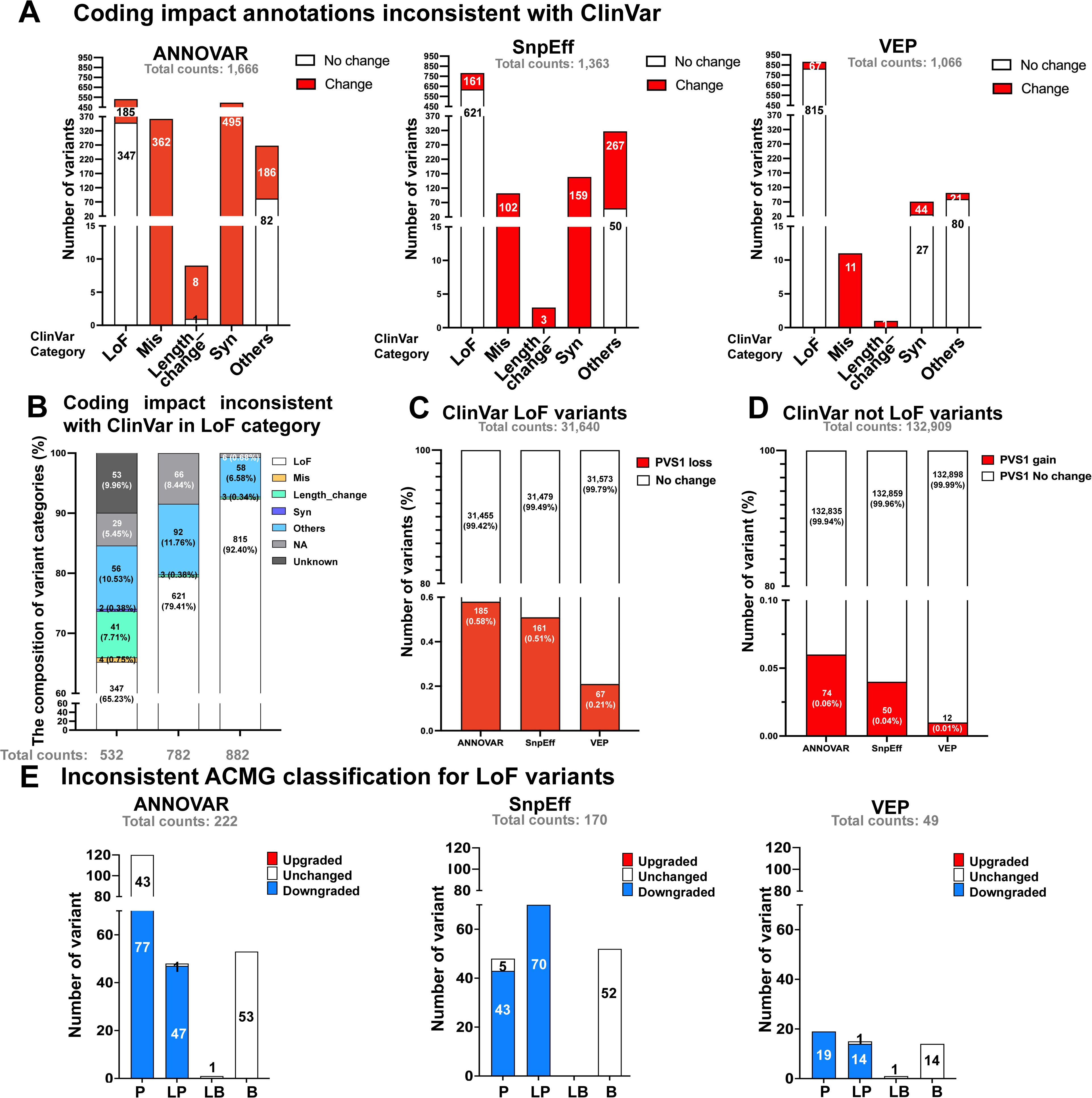
Impact of Discrepant Coding Impact Annotations on ACMG Classification. **(A)** Bar graph showing the total number of coding impact annotations inconsistent with ClinVar across the three tools (ANNOVAR, SnpEff, VEP), categorized by ClinVar functional consequence. Bars are divided into red (annotations resulting in a change in functional category) and white (annotations remaining unchanged) segments. Functional consequences include: LoF: loss of function; Mis: missense; Length_change: amino-acid length change; Syn: synonymous; Others. **(B)** Stacked bar chart to show the distribution of category changes for ClinVar-defined LoF variants that were misannotated by the tools. **(C)** Stacked bar graph to depict the number and proportion of ClinVar-defined LoF variants that lost PVS1 criterion due to misannotations by the tools. **(D)** Stacked bar graph to show the number and proportion of ClinVar-defined non-LoF variants (N=132,909) that gained the PVS1 criterion as a result of misannotations by the tools. **(E)** Bar graph comparing inconsistent ACMG classifications resulting from misannotations with bars representing the number of variants classified as Pathogenic (P), Likely Pathogenic (LP), Likely Benign (LB), and Benign (B) according to ClinVar. Each bar is divided into red (upgraded), white(unchanged) and blue (downgraded) segments based on curated classifications aligned with ACMG guidelines and ClinGen recommendations. The analyses focused on the impact of discrepant coding impact annotations on ACMG pathogenicity classifications. Inconsistencies in assigning the PVS1 criterion, which carries the heaviest weight in the ACMG guidelines, were found to significantly influence final variant interpretations, particularly for variants initially classified as pathogenic, likely pathogenic, or uncertain significance in ClinVar.

The composition of misannotated variants and the changes in annotation are illustrated in Figure 5B. ANNOVAR misannotated 532 LoF variants, leading to 185 variants changing categories while 347 remained unchanged (Figure 5A, 185/532=34.7%). Although VEP misannotated the highest number of LoF variants (N = 882), the majority of these (815/882 = 92.4%) did not change in their coding impact classification. Given that the LoF category represents the most heavily weighted criterion for the pathogenicity rules (PVS1), any inconsistencies may significantly influence the final interpretation of variants according to ACMG guidelines. For ClinVar-defined variants (LoF=31,640, not LoF=132,909), the analyses revealed that ANNOVAR accounted for the highest number of variants with PVS1 changes (PVS1-loss: 185, PVS1-gain: 74), followed by SnpEff (PVS1-loss: 161, PVS1-gain: 50), and VEP (PVS1-loss: 67, PVS1-gain: 12) (Figure 5C, 5D). To assess how functional category changes affect ACMG classification, we reevaluated all ACMG criteria for the variants with discrepant PVS1 changes to check whether curated ACMG classes aligned with ClinVar records. Automatic reevaluation was based on the comprehensive annotation using GDK platform. The rules PP3 has been disabled in the presence of PVS1. ClinVar lacks comprehensive details for applying ACMG rules necessitated excluding criteria like PS4, PM3, PP4, BP2, and BP5, which need gene-disease data, cohorts, or phasing for automated classification. Therefore, not all discrepant PVS1 variants were included in the reassessment. Figure 5E shows the results of the reassessment between ClinVar classification and the classification based on the corresponding annotation (ANNOVAR:222, SnpEff:170, VEP:49). This effect was particularly notable for PLP variants. Variants initially classified as BLB in ClinVar were less impacted by changes in PVS1. Incorrect interpretations of PVS1 led to substantial downgrades for PLP variants, with ANNOVAR downgrading 55.9% (124/222), SnpEff 66.5% (113/170), and VEP 67.3% (33/49), thereby increasing the risk of overlooking clinically relevant variants in reports. Our results indicate that inconsistencies in coding impact annotations, leading to discrepancies in PVS1 interpretation, significantly influenced final pathogenicity assessments.

## Discussion

Accurate variant annotation is essential for effective clinical diagnostics. We utilized ClinVar dataset to comprehensively compare the annotation capabilities of ANNOVAR, SnpEff, and VEP. In this study, we examined how transcript sets and annotation tools influence variant interpretation and their impact on ACMG classification. This study demonstrates the discrepancies among annotation tools can disconcert variant interpretation. Variability in transcript selection and HGVS nomenclature complicates variant retrieval and increases the risk of oversight during clinical genetic testing. Restricting transcripts to those selected by ClinVar improved consistency in HGVSc, HGVSp, and coding impact annotations; however, discrepancies with ClinVar persisted. Some HGVS syntax in ClinVar does not adhere to HGVS recommendations (see Table S3 and Table S4), and inconsistencies were between ClinVar web pages and its VCF. Moreover, gene inheritance modes provided by CGD^30^ ClinGen^31^, and PanelApp^32^ may also be inconsistent, impact the interpretation of PM2 and PM3 rules^5^ (Additional file 2: supplementary Figure S3). To mitigate these discrepancies, several strategies can be implemented.

1. **Standardize Annotation Practices:** Ensure uniformity in input by recalibrating data formats to utilize identical reference genomes and transcript annotations across all tools. The MANE transcripts should be broadly implemented. According to HGVS recommendations, “two variants separated by one nucleotide that together affect one amino acid should be described as a ‘delins.’” Multi-nucleotide variants (MNVs) must be phased and normalized in the VCF. HGVS syntax can complicate rules PS1/PM5, PS3/BS3, and PP5/BP6, underscoring the challenge for information retrieval. Improving the clarity of HGVS nomenclature is critical.
2. **Utilize Comprehensive Options**: Leverage MAF data from larger cohorts to enhance annotation accuracy. Utilize a full range of annotation options; for instance, using plug-in function --per_gene and --coding_only in VEP can streamline relevant transcripts, reducing complexity and potential errors. VEP “LOEUF”^33^ and “pLI”^34^ functions provide valuable insights into implicating gene LoF mechanisms. “NMD” plugin helps to predict if a variant’s position might be in a region that can escape nonsense-mediated decay (NMD)^35^. VEP’s “SameCodon” plugin helps for rule PS1 and PM5. “SingleLetterAA” for returning a HGVSp string with single amino acid letter codes. Prioritize high-impact variants, as they are often more clinically relevant.
3. **Cross-Validation and Review Protocols:** Implement cross-validation by comparing outputs from multiple independent tools for the same variants to identify discrepancies and determine the most accurate annotations. Use an in-house benchmark or ClinVar high-confidence set. Establish a protocol for regularly reviewing and updating annotation processes based on the latest research and tool capabilities, and ensure version control for each tool to facilitate data reanalysis.

## Supporting information

Key_resource

## Abbreviations

ACMG: American College of Medical Genetics and Genomics
AF: Allele frequency
AMP: American Association of Molecular Pathology
BLB: Benign and Likely Benign
Del: Deletion
Dup: Duplication
GDK: Gendiseak platform
HGVS: Human Genome Variation Society
HGVSc: The HGVS coding sequence name
HGVSp: The HGVS protein sequence name
Indel: Insertion-deletion
Inv: Inversion
Ins: Insertion
LOEUF: The loss-of-function observed/expected upper bound fraction
LoF: Loss-of-Function
MAF: Minor allele frequency
MANE: Matched Annotation from the NCBI and EMBL-EBI
MC: Molecular consequence
MS: Microsatellite
MT: Mitochondria
MNV: Multi Nucleotide Variants
NGS: Next-Generation Sequencing
NMD: Nonsense-Mediated Decay
Nc: Non-coding
PLP: Pathogenic and Likely Pathogenic
SO: Sequence Ontology
SNV: Single Nucleotide Variant
VCF: Variant Call Format File
VEP: Ensembl Variant Effect Predictor
VUS: Variant of Uncertain Significance

## Declarations

### Data Availability Statement

All data relevant to this study have been submitted along with the manuscript to the journal. The code used in this study has been uploaded to GitHub and is accessible at https://github.com/YuanTzuH/AnnotatorCompare.

## Acknowledgments

We appreciate the technical assistance of Taiwan AI labs. We are also grateful to the National Center for High-performance Computing (NCHC) for providing computational and storage resources. This study was supported by research grants from the National Science and Technology Council in Taiwan (NSTC 111-2320-B-002-091-MY3, 112-2320-B-002-045, 112-2622-B-002-012) and National Taiwan University (113L892901).

## Author Contributions

Conceptualization: Yu-An Chen, Jia-Hsing Huang, Jacob Shu-Jui Hsu; Data curation: Yu-An Chen, Tzu-Hang Yuan; Formal analysis: Yu-An Chen, Tzu-Hang Yuan; Funding acquisition: Jacob Shu-Jui Hsu; Methodology: Yu-An Chen, Tzu-Hang Yuan; Software: Tzu-Hang Yuan, Jia-Hsing Huang, Yu-Bin Wang, Tzu-Mao Hung; Supervision: Jacob Shu-Jui Hsu, Pei-Lung Chen, Chien-Yu Chen; Validation: Yu-An Chen, Jacob Shu-Jui Hsu; Visualization: Yu-An Chen, Tzu-Hang Yuan; Writing-original draft: Yu-An Chen; Writing-review & editing: Jacob Shu-Jui Hsu, Pei-Lung Chen, and all authors.

## Ethics Declaration

This study did not involve the collection of human or animal samples, and therefore, Institutional Review Board (IRB) approval was not required.

## Declaration of AI and AI-assisted technologies in the writing process

During the preparation of this work, the author(s) used Perplexity AI, a natural language model trained on a diverse dataset up to October 2023, in order to enhance the coherence and overall readability. After using this tool/service, the author(s) reviewed and edited the content as needed and take(s) full responsibility for the content of the publication.

## Key Resource Table

Software and dataset used in this study.

## Supplementary Materials

### Additional file 1: supplementary tables

**Table S1.** Consequence prioritization and SO term normalization.

**Table S2.** The Annotation information and corresponding ACMG rules are discussed in this study.

**Table S3.** Exemplar variants demonstrating HGVSc nomenclature discrepancies.

**Table S4.** Exemplar variants demonstrating HGVSp nomenclature discrepancies.

### Additional file 2:supplemtntary figures

**Figure S1. Methodology of HGVS syntax comparison.** We applied the following assessments to compare two HGVS expressions in our dataset. The query transcript or gene symbol must match the reference. **(A)** For the HGVSc comparison, if the accession does not match, the variant is not assessed for the following matching. **(B)** For HGVSp comparison, if the gene symbol corresponding to the transcript does not match, the variant is not evaluated for the following matching. An exact match and an equivalent match of syntax are considered matches. If the syntax is not an alternative expression of the other HGVS variant, the match is considered “incorrect.”

**Figure S2. Discordant HGVS with ClinVar across tools. (A)** The distribution of causes of discordant HGVSc. **(B)** The distribution of causes of discordant HGVSp. NA indicate missing HGVS or information could not be retrieved.

**Figure S3. Mode of inheritance (MOI) information obtained from databases**. AD: autosomal dominant. AR: autosomal recessive. XL: X-linked. YL: Y-linked.

**Additional file 3. Precedure of data process and evaluation.**

**Additional file 4. Fig4_supplementary data**

**Additional file 5. Fig5_supplementary data**

**Additional file 6. Fig5E_supplementary_ANNOVAR_PVS1_change**

**Additional file 7. Fig5E_supplementary_SnpEff_PVS1_change**

**Additional file 8. Fig5E_supplementary_VEP_PVS1_change**

