## Supplementary_tables for "Toward Automatic Variant Interpretation: Discordant Genetic Interpretation Across Variant Annotations for ClinVar Pathogenic Variants"

**Table S1. Consequence prioritization and SO term normalization. The terms in the table below are shown in order of severity (more severe to less severe).**

| IMPACT | | SO | ClinVar | ANNOVAR | SnpEff | VEP |
|---|---|---|---|---|---|---|
| HIGH | High | **splicing** | splice_acceptor | splicing | splice_acceptor | splice_acceptor |
| | | | splice_donor | . | splice_donor | splice_donor |
| | High | **nonsense** | nonsense | stopgain | stop_gained | stop_gained |
| | High | **frameshift** | frameshift | frameshift deletion | frameshift | frameshift |
| | | | . | frameshift insertion | . | . |
| | High | **stop_lost** | | stoploss | stop_lost | stop_lost |
| | High | **initiatior_codon** | initiatior_codon | . | initiator_codon | . |
| | | | . | startloss | start_lost | start_lost |
| | Moderate | **inframe** | inframe_deletion | nonframeshift deletion | disruptive_inframe_deletion | inframe_deletion |
| | | | inframe_insertion | nonframeshift insertion | disruptive_inframe_insertion | Inframe_insertion |
| | | | inframe_indel | . | conservative_inframe_deletion | . |
| | | | | . | conservative_inframe_insertion | . |
| | Moderate | **missense** | missense | nonsynonymous SNV | missense | missense |
| | | | . | nonframeshift substitution | . | . |
| | Moderate | **synonymous** | synonymous | synonymous SNV | synonymous | synonymous |
| | | | . | . | stop_retained | . |
| | Low | **5_prime_UTR** | 5_prime_UTR | UTR5 | 5_prime_UTR | 5_prime_UTR |
| | Low | **3_prime_UTR** | 3_prime_UTR | UTR3 | 3_prime_UTR | 3_prime_UTR |
| | Low | **non_coding_transcript** | non-coding_transcript | ncRNA_exonic | non_coding_transcript_exon | non_coding_transcript_exon |
| | | | . | ncRNA_intronic | non_coding_transcript | non_coding_transcript |
| | | | . | ncRNA_splicing | . | . |
| | Low | **intron** | intron | intronic | intron | intron |
| | | | . | . | . | splice_donor_5th_base |
| | | | . | . | splice_region | splice_region |
| | | | . | . | . | splice_donor_region |
| | | | . | . | . | splice_polypyrimidine_tract |
| | Low | **genic_upstream_transcript** | genic_upstream_transcript | upstream | upstream_gene | upstream_gene |
| | Low | **genic_downstream_transcript** | genic_downstream_transcript | downstream | downstream_gene | downstream_gene |
| | | **intergenic** | intergenic | intergenic | intergenic | intergenic |
| LOW | | | . | unknown | . | . |

**IMPACT: HIGH**
The variant is assumed to have high (disruptive) impact in protein, probably causing protein truncation, loss of function or triggering nonsense mediated decay.

**IMPACT: MODERATE**
A non-disruptive variant that might change protein effectiveness.

**IMPACT: LOW**
Assumed to be mostly harmless or unlikely to change protein behavior.

**Table S2. The Annotation information and corresponding ACMG rules discussed in this study.**

| Annotation information | | Corresponding ACMG rule | Details |
|---|---|---|---|
| HGVS | syntax-based retrieval | PS1 | PS1 Same amino acid change as a previously established pathogenic variant regardless of nucleotide change |
| | | PM5 | Novel missense change at an amino acid residue where a different missense change determined to be pathogenic has been seen before |
| | | PS3 | Well-established in vitro or in vivo functional studies supportive of a damaging effect on the gene or gene product |
| | | BS3 | Well-established in vitro or in vivo functional studies show no damaging effect on protein function or splicing |
| | | PP5 | Reputable source recently reports variant as pathogenic, but the evidence is not available to the laboratory to perform an independent evaluation |
| | | BP6 | Reputable source recently reports variant as benign, but the evidence is not available to the laboratory to perform an independent evaluation |
| Coding impact consequence | LoF | PVS1 | PVS1 null variant (nonsense, frameshift, canonical ±1 or 2 splice sites, initiation codon, single or multiexon deletion) in a gene where LOF is a known mechanism of disease |
| | missense | PM1 | Located in a mutational hot spot and/or critical and well-established functional domain (e.g., active site of an enzyme) without benign variation |
| | | PP2 | Missense variant in a gene that has a low rate of benign missense variation and in which missense variants are a common mechanism of disease |
| | | BP1 | Missense variant in a gene for which primarily truncating variants are known to cause disease |
| | protein length-change | PM4 | Protein length changes as a result of in-frame deletions/insertions in a nonrepeat region or stop-loss variants |
| | | BP3 | In-frame deletions/insertions in a repetitive region without a known function |
| | synonymous | BP7 | A synonymous (silent) variant for which splicing prediction algorithms predict no impact to the splice consensus sequence nor the creation of a new splice site AND the nucleotide is not highly conserved |

### Table S3. Exemplar variants demonstrating HGVSc nomenclature discrepancies.

| Variant type | ClinVar | ANNOVAR | SnpEff | VEP | HGVS preference (VariantValidator) |
|---|---|---|---|---|---|
| Substitution | c.351C>T | c.324C>T | c.351C>T | c.351C>T | NM_001378609.3:c.351C>T |
| Substitution | c.405C>T | c.426C>T | c.405C>T | c.405C>T | NM_001282684.2:c.405C>T |
| Substitution | c.1981G>A | c.1954G>A | c.1981G>A | c.1981G>A | NM_001378609.3:c.1981G>A |
| Substitution | c.4530G>C | c.4656G>C | c.4530G>C | c.4530G>C | NM_020884.7:c.4530G>C |
| Deletion | c.623_624del | c.618_619del | c.623_624delCC | c.623_624del | NM_001006658.3:c.623_624del |
| Deletion | c.2259_2260del | c.2258_2259del | c.2259_2260delAA | c.2259_2260del | NM_001382391.1:c.2259_2260del |
| Deletion | c.2348_2351del | c.2347_2350del | c.2348_2351delCGCA | c.2348_2351del | NM_006767.4:c.2348_2351del |
| Deletion | c.1053del | c.1049delA | c.1053delA | c.1053del | NM_000059.4:c.1053del |
| Duplication | c.1908_1909dup | c.1906_1907insGT | c.1908_1909dupTG | c.1908_1909dup | NM_000038.6:c.1908_1909dup |
| Duplication | c.819dup | c.819dupT | c.819dupT | c.819_820insT | NM_024577.4:c.819dup |
| Duplication | c.243dup | c.241dupA | c.243dupA | c.243dup | NM_001134673:c.243dup |
| Duplication | c.667_674dup | c.674_675insAACATTCC | c.667_674dupAACATTCC | c.674_675insAACATTCC | NM_000277.3:c.667_674dup |
| Insersion | c.6449_6450insTA | c.6448_6449insAT | c.6449_6450insTA | c.6449_6450insTA | NM_000059.4:c.6449_6450insTA |
| Insersion | c.36_37insCG | c.35_36insGC | c.36_37insCG | c.36_37insCG | NM_000642.3:c.36_37insCG |
| Insersion | c.433_434insGCTGTTA | c.432_433insAGCTGTT | c.433_434insGCTGTTA | c.433_434insGCTGTTA | NM_000532.5:c.433_434insGCTGTTA |
| Insersion | c.1487_1488insGGCG | c.1485_1486insCGGG | c.1487_1488insGGCG | c.1487_1488insGGCG | NM_000112.4:c.1487_1488insGGCG |
| Inversion | c.773_774inv | c.773_774delinsCA | c.773_774delTGinsCA | c.773_774inv | NM_015443.4:c.773_774inv |
| Inversion | c.47_48inv | c.47_48delinsCA | c.47_48delTGinsCA | c.47_48inv | NM_000260.4:c.47_48inv |
| Inversion | c.1719_1720inv | c.1719_1720delinsTG | c.1719_1720delCAinsTG | c.1719_1720inv | NM_000545.8:c.1719_1720inv |
| Inversion | c.5843_5844inv | c.5843_5844delinsTG | c.5843_5844delCAinsTG | c.5843_5844inv | NM_000350.3:c.5843_5844inv |
| Repeat | c.4213AAG[3] | c.4222_4224del | c.4222_4224delAAG | c.4222_4224del | NM_000552.5:c.4222_4224del |
| Repeat | c.337GCGCCCGCC[3] | c.354_355insGCGCCCGCC | c.346_354dupGCGCCCGCC | c.354_355insGCGCCCGCC | NM_001135649.3:c.336_337insCGCGCCCGC |
| Repeat | c.66GTCGCC[3] | c.84_89del | c.84_89delGTCGCC | c.84_89del | NM_001018111.3:c.84_89del |
| Repeat | c.3528CAG[10] | c.3545_3546insCAGCAGCAGCAG | c.3534_3545dupCAGCAGCAGCAG | c.3545_3546insCAGCAGCAGCAG | NM_016580.4:c.3527_3528insGCAGCAGCAGCA |

### Table S4. Exemplar variants demonstrating HGVSp nomenclature discrepancies.

| Variant type | ClinVar | ANNOVAR | SnpEff | VEP | HGVS preference (VariantValidator) |
|---|---|---|---|---|---|
| Synonymous | p.Asn117= | p.N108N | p.Asn117Asn | p.Asn117= | NP_001365538.2:p.(Asn117=) |
| Synonymous | p.Ala817= | p.A859A | p.Ala817Ala | p.Ala817= | NP_065935.4:p.(Ala817=) |
| Synonymous | p.His135= | p.H142H | p.His135His | p.His135= | NP_001269613.2:p.(His135=) |
| Synonymous | p.Phe516= | p.F558F | p.Phe516Phe | p.Phe516= | NP_065935.4:p.(Phe516=) |
| Missense | p.Ala19Thr | p.A26T | p.Ala19Thr | p.Ala19Thr | NP_001269613.2:p.(Ala19Thr) |
| Missense | p.Ser200Ile | p.S184I | p.Ser200Ile | p.Ser200Ile | NP_001094891.4:p.(Ser200Ile) |
| Missense | p.Cys653Gly | p.C644G | p.Cys653Gly | p.Cys653Gly | NP_001365538.2:p.(Cys653Gly) |
| Nonsense | p.Arg360Ter | p.R351X | p.Arg360* | p.Arg360Ter | NP_001365538.2:p.(Arg360Ter) |
| Nonsense | p.Gln529Ter | p.Q520X | p.Gln529* | p.Gln529Ter | NP_001365538.2:p.(Gln529Ter) |
| Nonsense | p.Cys3073_Asp3074delinsTer | p.C3073* | p.Cys3073fs | p.Cys3073Ter | NP_055178.3:p.(Cys3073Ter) |
| Nonsense | p.Ser160_Ser161insTer | p.S161* | p.Ser161fs | p.Ser161Ter | NP_892006.3:p.(Ser161Ter) |
| Frameshift | p.Asp415fs | p.D415Efs*20 | p.Asp415fs | p.Asp415GlufsTer20 | NP_005350.1:p.(Asp415GlufsTer20) |
| Frameshift | p.Met128fs | p.L129Gfs*14 | p.Met128fs | p.Met128ValfsTer11 | NP_510965.1:p.(Met128ValfsTer11) |
| Frameshift | p.Thr1410Asnfs | p.T1410Nfs*4 | p.Thr1410fs | p.Thr1410AsnfsTer4 | NP_000050.2:p.(Thr1410AsnfsTer4) |
| Frameshift | p.Ile1036Hisfs | p.I1036Hfs*14 | p.Ile1036fs | p.Ile1036HisfsTer14 | NP_001124459.1:p.(Ile1036HisfsTer14) |
| Extension | p.Ter453ValextTer? | p.*453delinsVKPWTECGLSAVNLLMEI QLFLSSEKVRKANLNLK* | p.Ter453fs | p.Ter453ValfsTer36 | NP_000268.1:p.(Ter453ValextTer35) |
| Extension | p.Ter380Arg | p.X380R | p.Ter380Argext*? | p.Ter380ArgextTer49 | NP_000146.2:p.(Ter380ArgextTer49) |
| Extension | p.Ter499Trp | p.X499W | p.Ter499Trpext*? | p.Ter499TrpextTer27 | NP_001104262.1:p.(Ter499TrpextTer27) |
| Extension | p.Ter741Cys | p.X741C | p.Ter741Cysext*? | p.Ter741CysextTer13 | NP_659475.2:p.(Ter741CysextTer13) |
| Inframe | p.Arg80_Leu82del | p.R80_L82del | p.Leu79_Leu81del | p.Arg80_Leu82del | NP_000024.2:p.(Arg80_Leu82del) |
| Inframe | p.Thr51_Thr54del | p.T51_T54del | p.Thr50_Val53del | p.Thr51_Thr54del | NP_001138498.1:p.(Thr51_Thr54del) |
| Inframe | p.Ile112dup | p.I112_A113insI | p.Ile112dup | p.Ile112dup | NP_000093.1:p.(Ile112dup) |
| Inframe | p.Cys169_Ser173dup | p.S173_W174insCNLTS | p.Cys169_Ser173dup | p.Cys169_Ser173dup | NP_000147.1:p.(Cys169_Ser173dup) |
