## Supplementary_figures for "Toward Automatic Variant Interpretation: Discordant Genetic Interpretation Across Variant Annotations for ClinVar Pathogenic Variants"

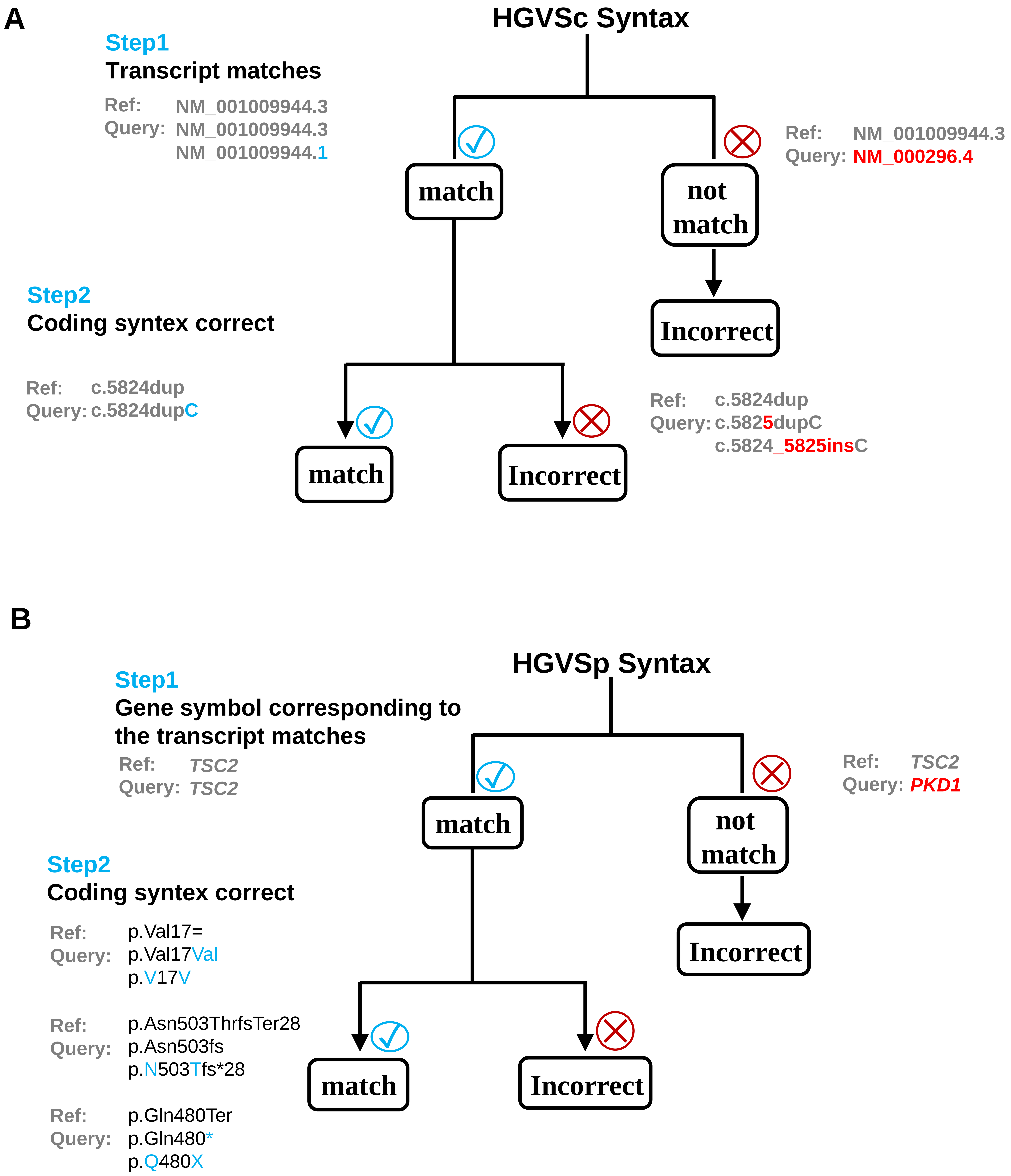


**Figure S1. Methodology of HGVS syntax comparison.** To compare two HGVS expressions in our dataset, we applied the following assessments. The query transcript or gene symbol must match the reference. **(A)** For HGVSc comparison, if the accession does not match, the variant is not assessed the following matching. **(B)** For HGVSp comparison, if the gene symbol corresponding to the transcript does not match, the variant is not assessed the following matching. Both exact match and equivalent match of the syntax are considered as a match. If the syntax is not an alternative expression of the other HGVS variant, the match is considered "incorrect."

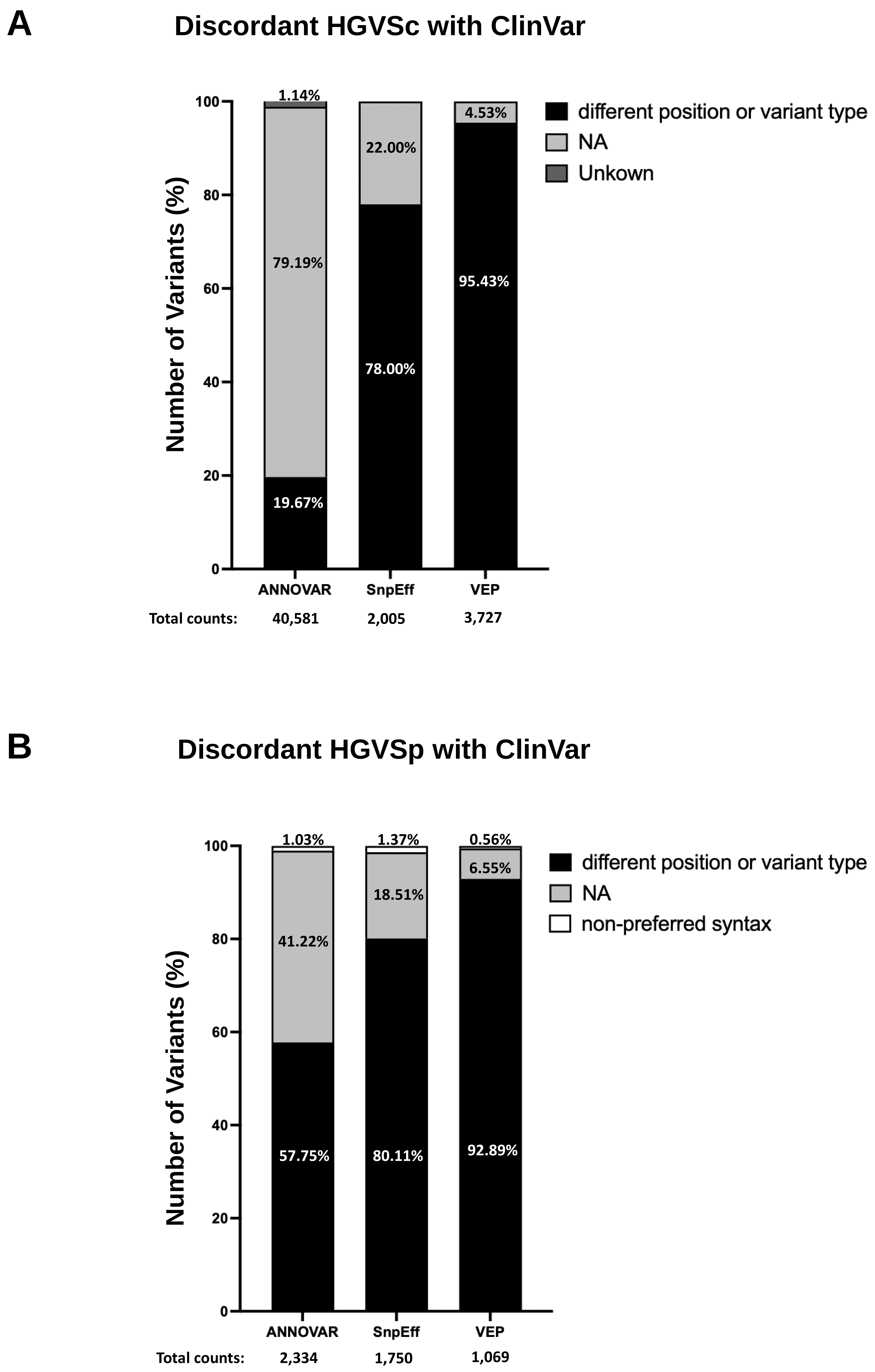


**Figure S2. Discordant HGVS with ClinVar across tools. (A)** The distribution of causes of discordant HGVSc. **(B)** The distribution of causes of discordant HGVSp. NA indicate missing HGVS or information could not been retrieved.

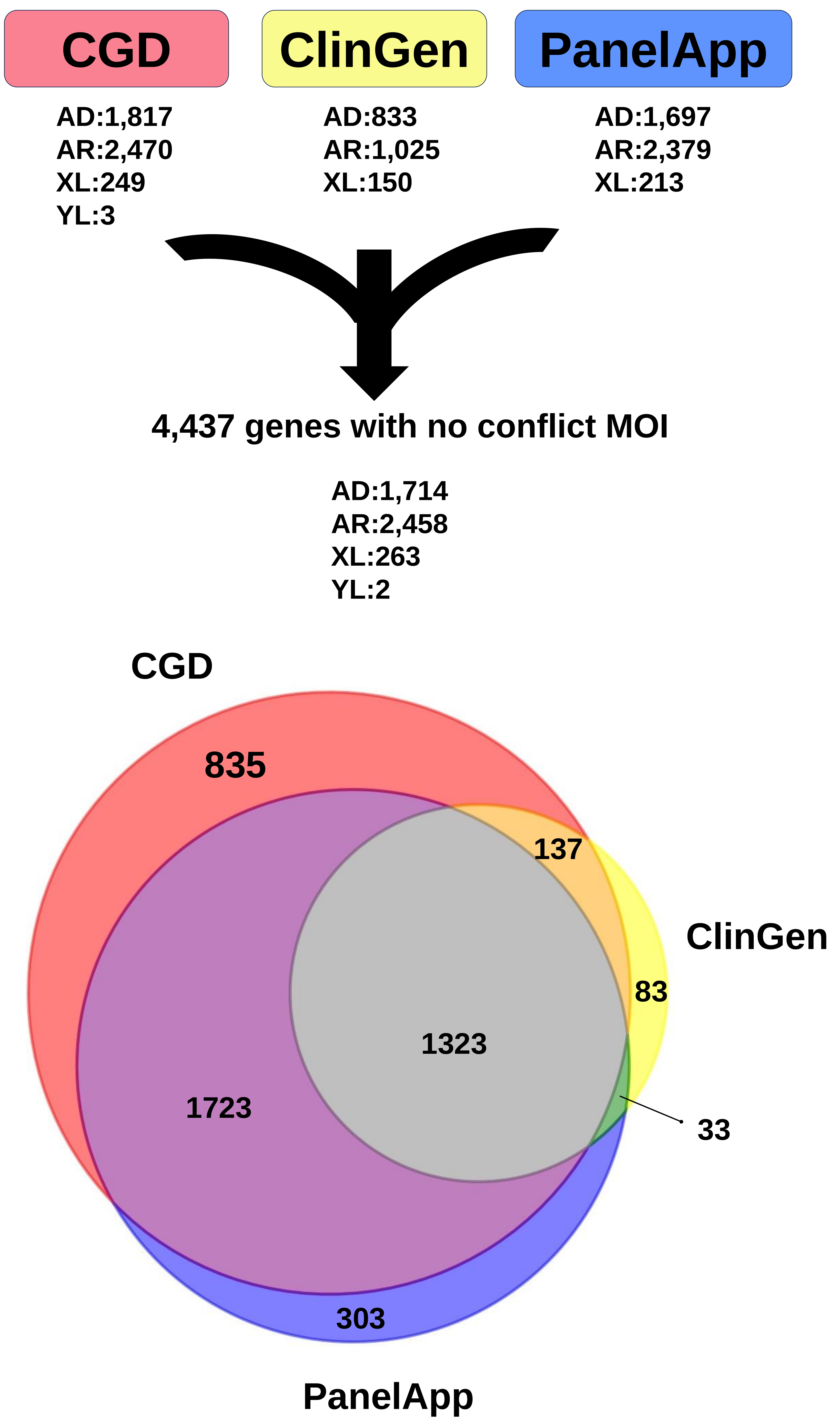


**Figure S3. Mode of inheritance (MOI) information obtained from databases**. AD: autosomal dominant. AR: autosomal recessive. XL:X-linked. YL:Y-linked.
