## Supplementary_file for "Toward Automatic Variant Interpretation: Discordant Genetic Interpretation Across Variant Annotations for ClinVar Pathogenic Variants"

### Precedure of data process and evaluation

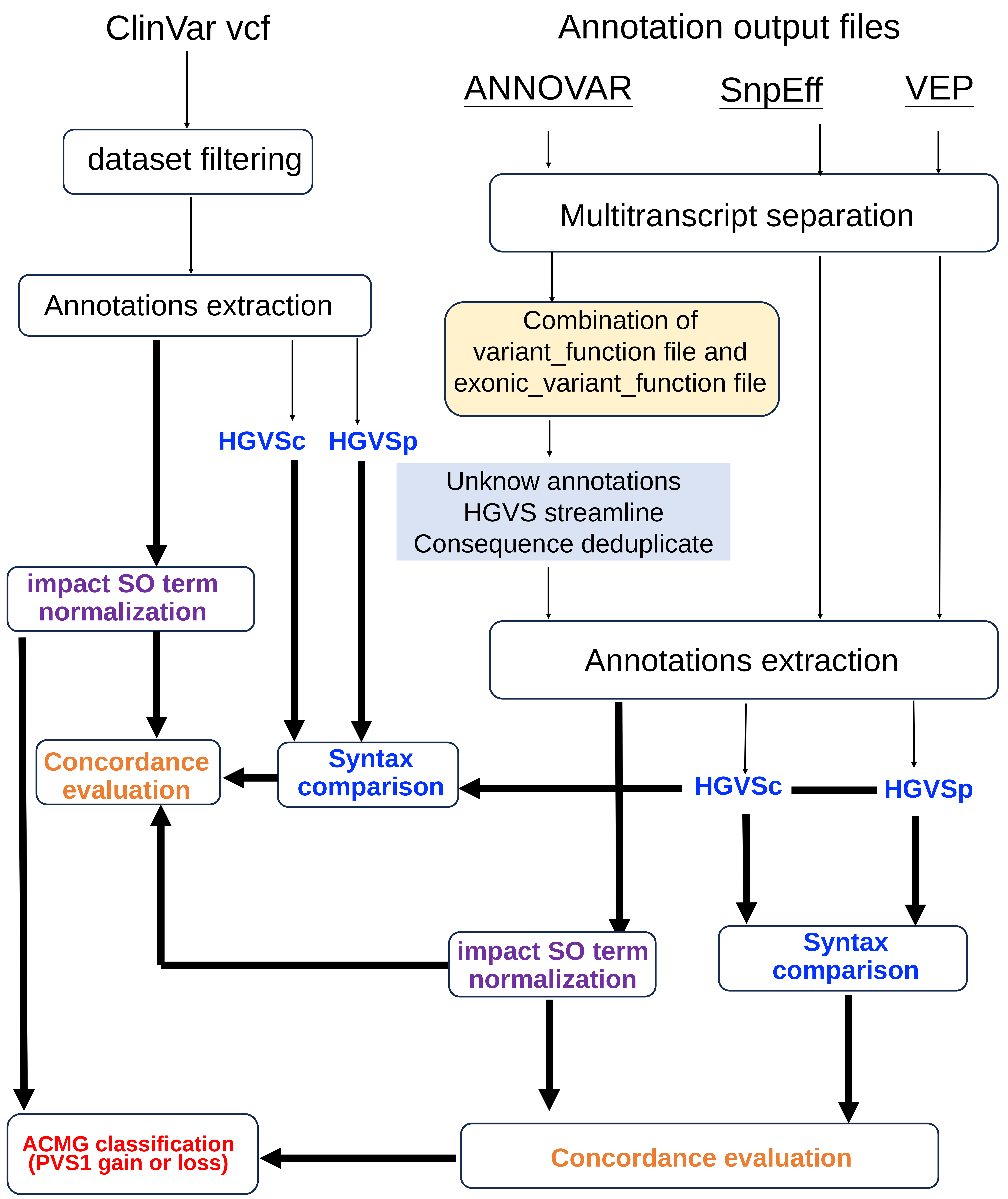

### ANNOVAR annotation output file process (PLP variants)

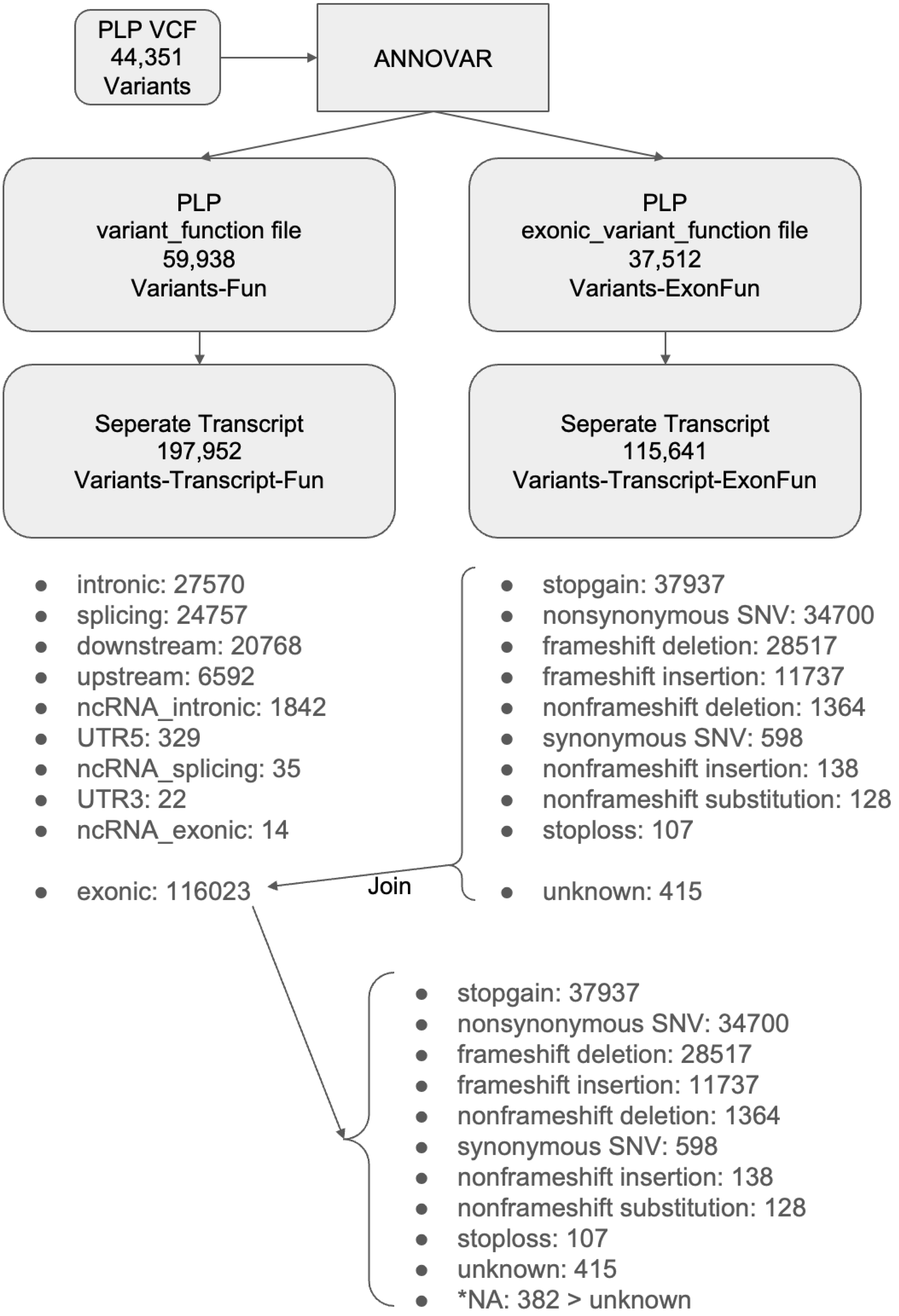

### ANNOVAR annotation output file process (BLB variants)

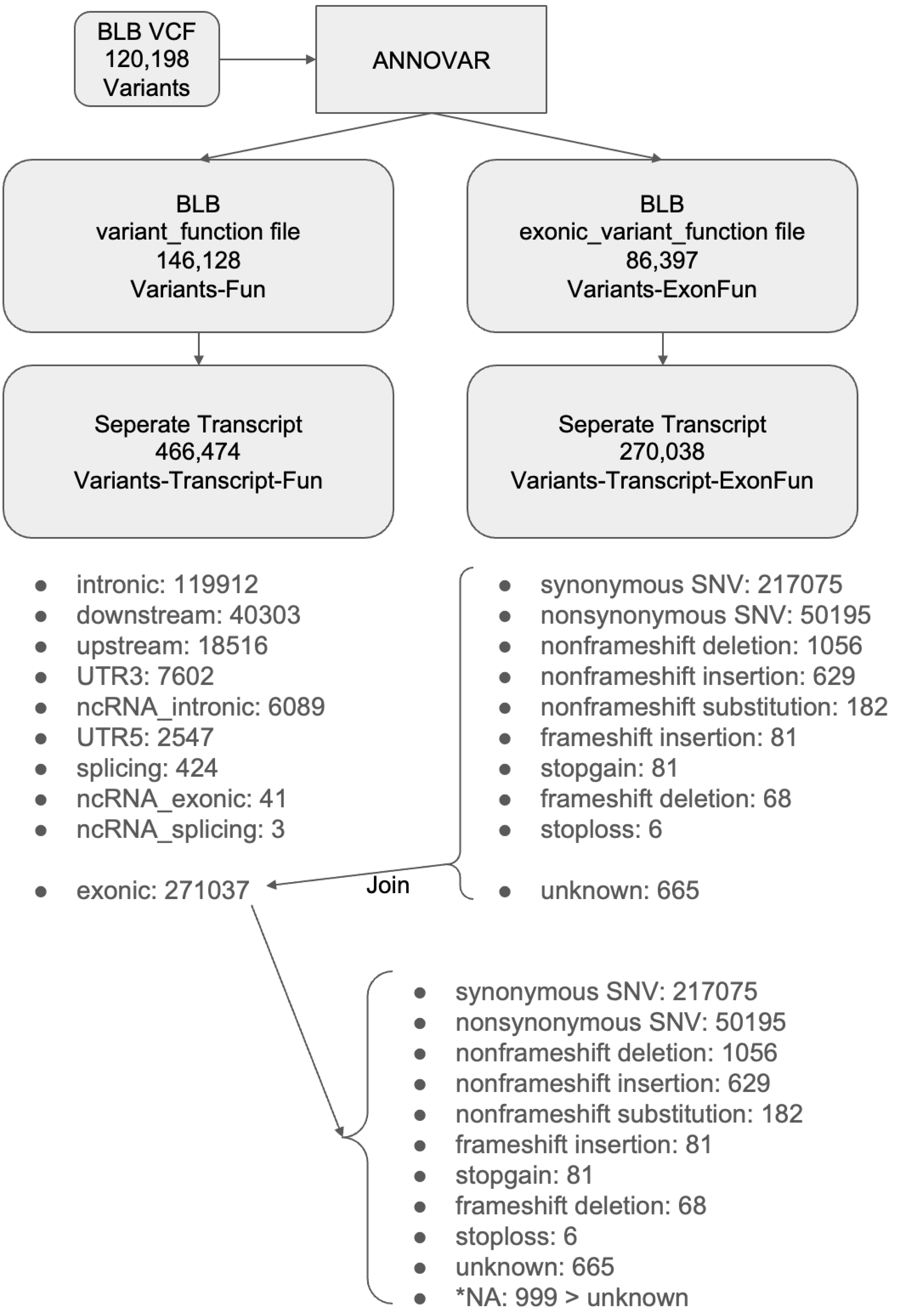

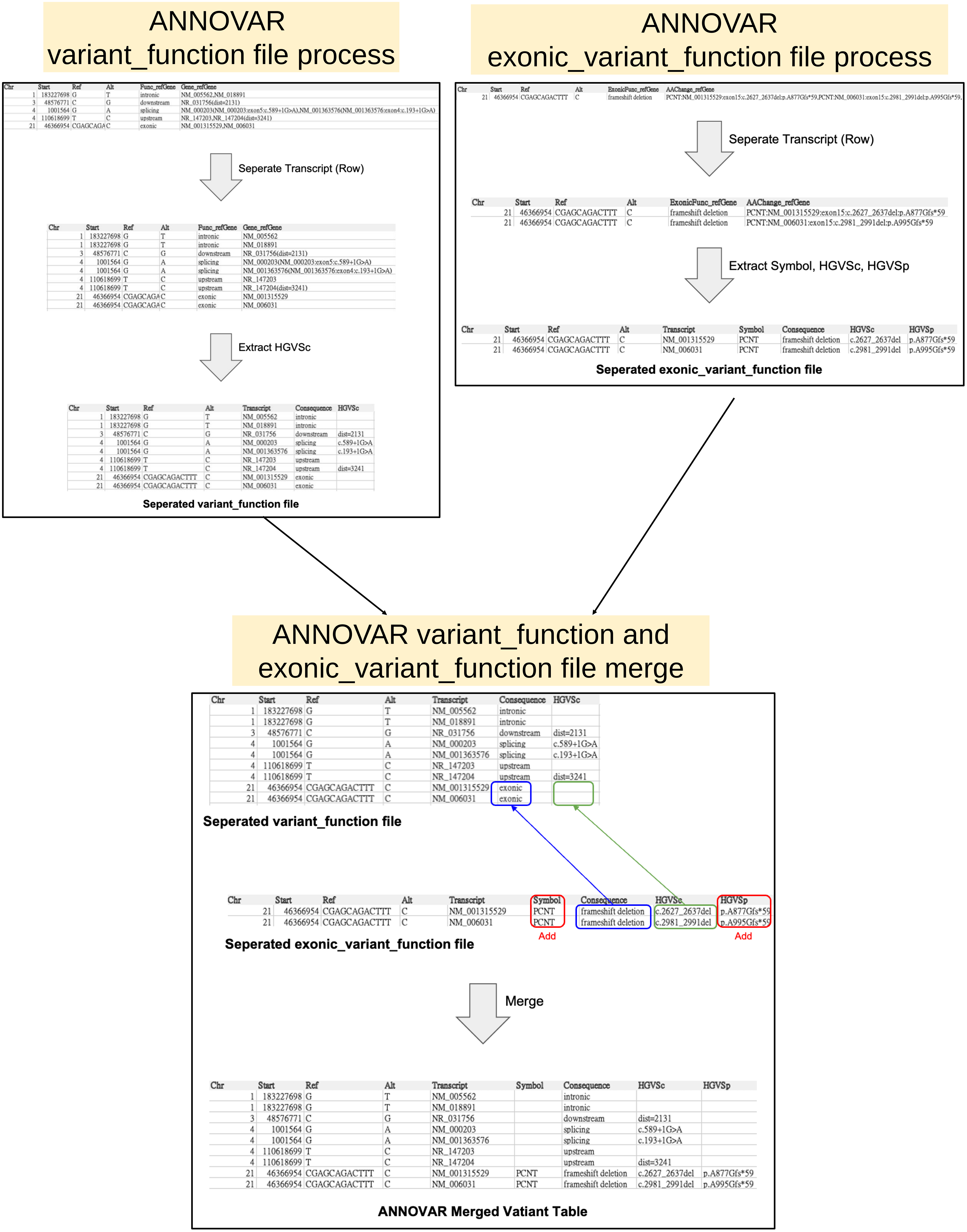
ANNOVAR
variant_function file process
ANNOVAR
exonic_variant_function file process
Seperate Transcript (Row)
Extract HGVSc
Seperated variant_function file
Seperate Transcript (Row)
Extract Symbol, HGVSc, HGVSp
Seperated exonic_variant_function file
ANNOVAR variant_function and
exonic_variant_function file merge
Seperated variant_function file
Seperated exonic_variant_function file
Add
Add
Merge
ANNOVAR Merged Vatiant Table

### ANNOVAR unknown annotation

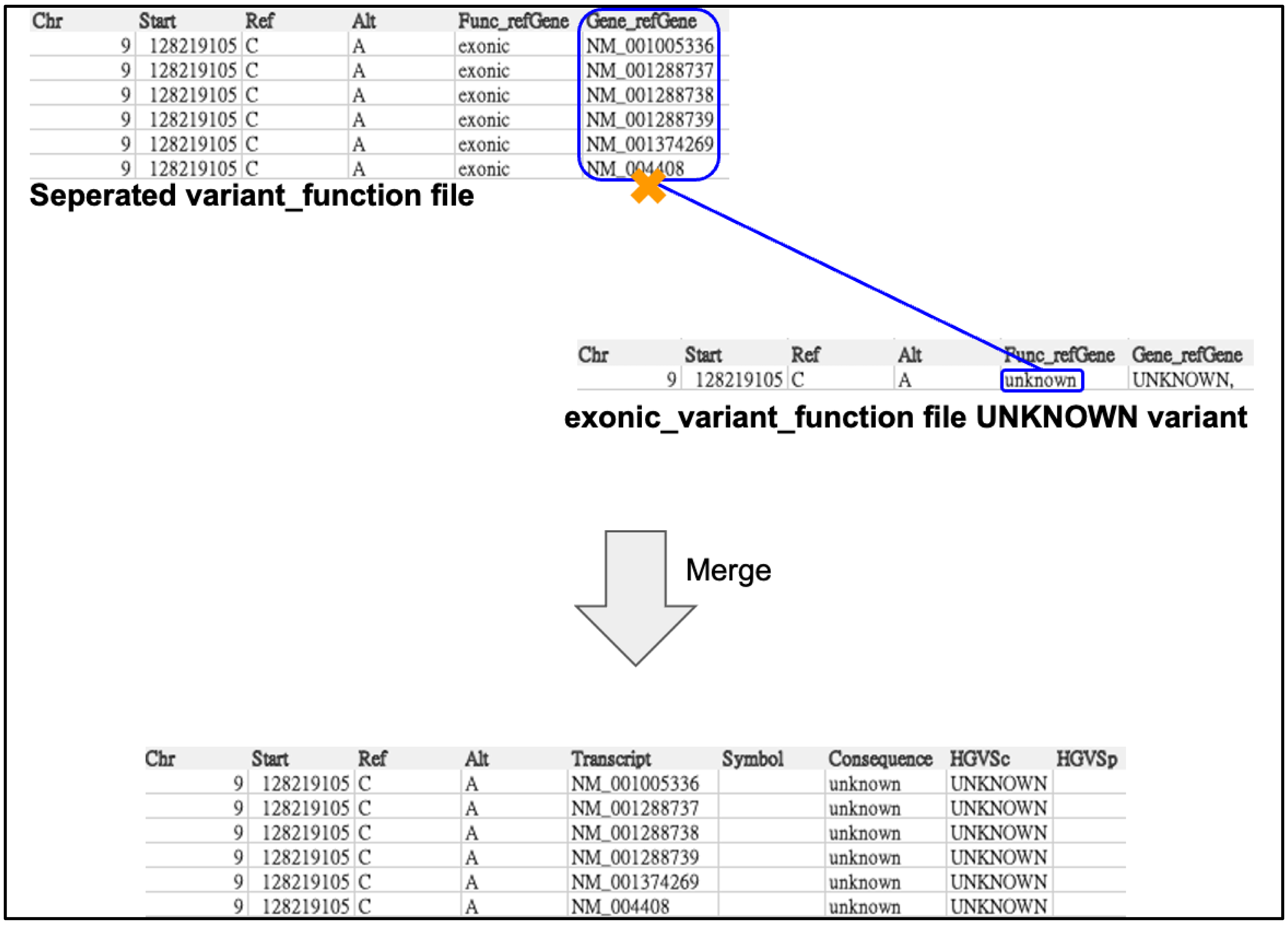


| Chr | Start | Ref | Alt | Func_refGene | Gene_refGene |
|---|---|---|---|---|---|
| 9 | 128219105 | C | A | exonic | NM_001005336 |
| 9 | 128219105 | C | A | exonic | NM_001288737 |
| 9 | 128219105 | C | A | exonic | NM_001288738 |
| 9 | 128219105 | C | A | exonic | NM_001288739 |
| 9 | 128219105 | C | A | exonic | NM_001374269 |
| 9 | 128219105 | C | A | exonic | NM_004408 |

| Chr | Start | Ref | Alt | Func_refGene | Gene_refGene |
|---|---|---|---|---|---|
| 9 | 128219105 | C | A | unknown | UNKNOWN, |

| Chr | Start | Ref | Alt | Transcript | Symbol | Consequence | HGVSc | HGVSp |
|---|---|---|---|---|---|---|---|---|
| 9 | 128219105 | C | A | NM_001005336 | | unknown | UNKNOWN | |
| 9 | 128219105 | C | A | NM_001288737 | | unknown | UNKNOWN | |
| 9 | 128219105 | C | A | NM_001288738 | | unknown | UNKNOWN | |
| 9 | 128219105 | C | A | NM_001288739 | | unknown | UNKNOWN | |
| 9 | 128219105 | C | A | NM_001374269 | | unknown | UNKNOWN | |
| 9 | 128219105 | C | A | NM_004408 | | unknown | UNKNOWN | |

### ANNOVAR HGVS streamline

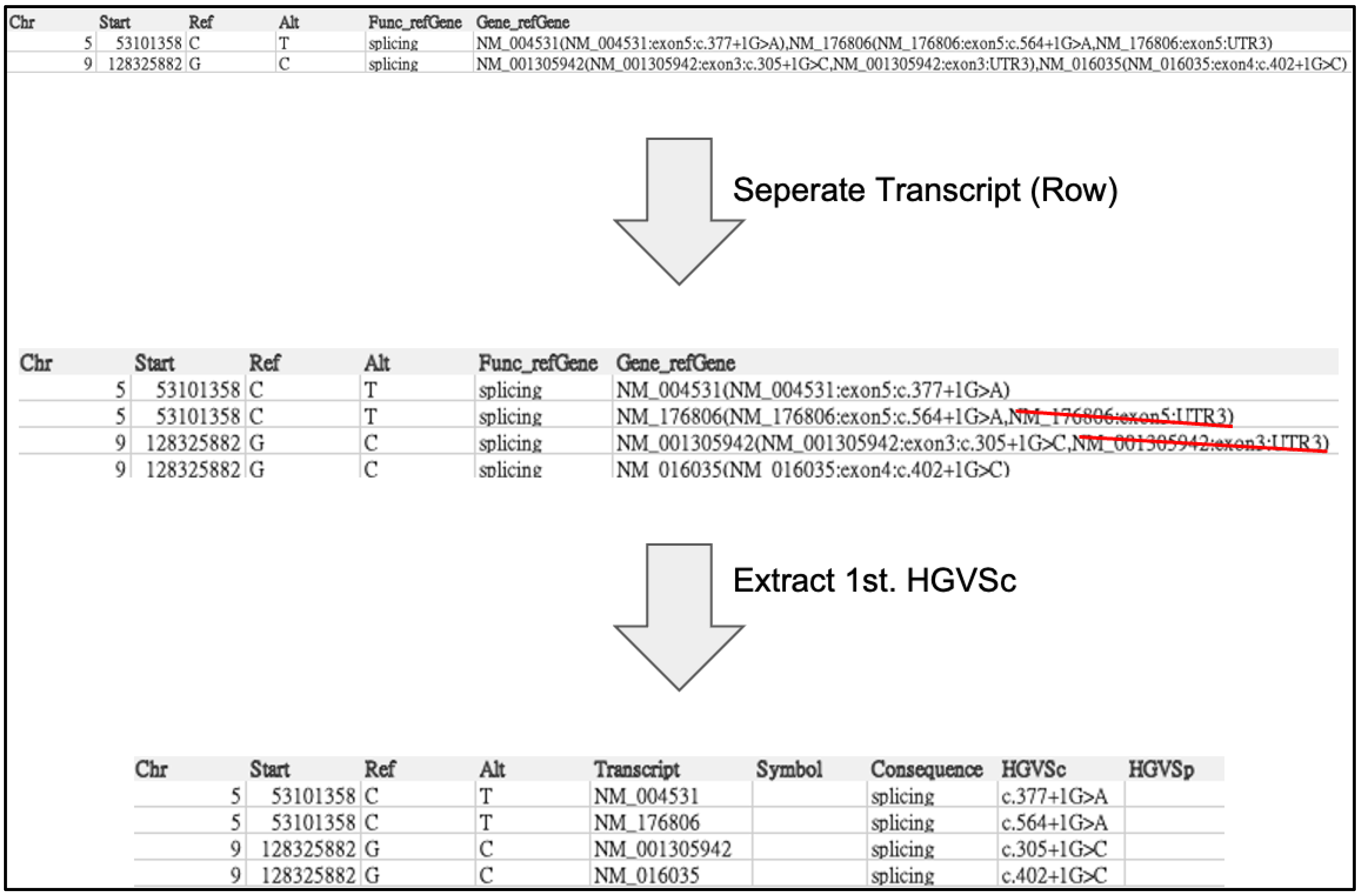


| Chr | Start | Ref | Alt | Func_refGene | Gene_refGene |
|---|---|---|---|---|---|
| 5 | 53101358 | C | T | splicing | NM_004531(NM_004531:exon5:c.377+1G>A),NM_176806(NM_176806:exon5:c.564+1G>A,NM_176806:exon5:UTR3) |
| 9 | 128325882 | G | C | splicing | NM_001305942(NM_001305942:exon3:c.305+1G>C,NM_001305942:exon3:UTR3),NM_016035(NM_016035:exon4:c.402+1G>C) |

| Chr | Start | Ref | Alt | Func_refGene | Gene_refGene |
|---|---|---|---|---|---|
| 5 | 53101358 | C | T | splicing | NM_004531(NM_004531:exon5:c.377+1G>A) |
| 5 | 53101358 | C | T | splicing | NM_176806(NM_176806:exon5:c.564+1G>A,NM_176806:exon5:UTR3) |
| 9 | 128325882 | G | C | splicing | NM_001305942(NM_001305942:exon3:c.305+1G>C,NM_001305942:exon3:UTR3) |
| 9 | 128325882 | G | C | splicing | NM_016035(NM_016035:exon4:c.402+1G>C) |

| Chr | Start | Ref | Alt | Transcript | Symbol | Consequence | HGVSc | HGVSp |
|---|---|---|---|---|---|---|---|---|
| 5 | 53101358 | C | T | NM_004531 | | splicing | c.377+1G>A | |
| 5 | 53101358 | C | T | NM_176806 | | splicing | c.564+1G>A | |
| 9 | 128325882 | G | C | NM_001305942 | | splicing | c.305+1G>C | |
| 9 | 128325882 | G | C | NM_016035 | | splicing | c.402+1G>C | |

### ANNOVAR consequence deduplicate

| key <chr> | chr <chr> | start <dbl> | ref <chr> | alt <chr> | NM <chr> | Consequence <chr> | HGVSc <chr> | Symbol <chr> | HGVSp <chr> |
|---|---|---|---|---|---|---|---|---|---|
| 167569\|178004 | 1 | 145926947 | G | T | NM_005105 | intronic | . | . | . |
| 167569\|178004 | 1 | 145926947 | G | T | NR_002328 | downstream | . | . | . |
| 167569\|178004 | 1 | 145926947 | G | T | NR_104033 | downstream | dist=1606 | . | . |
| 167569\|178004 | 1 | 145926947 | G | T | NR_147182 | ncRNA_intronic | . | . | . |
| 167569\|178004 | 1 | 145926947 | G | T | NR_147182 | ncRNA_splicing | c.116+1G>T | . | . |
| 2049534\|2107421 | 9 | 127818309 | A | G | NM_000118 | synonymous SNV | c.1497T>C | ENG | p.P499P |
| 2049534\|2107421 | 9 | 127818309 | A | G | NM_001018078 | downstream | . | . | . |
| 2049534\|2107421 | 9 | 127818309 | A | G | NM_001114753 | synonymous SNV | c.1497T>C | ENG | p.P499P |
| 2049534\|2107421 | 9 | 127818309 | A | G | NM_001278138 | synonymous SNV | c.951T>C | ENG | p.P317P |
| 2049534\|2107421 | 9 | 127818309 | A | G | NM_001288803 | downstream | . | . | . |
| 2049534\|2107421 | 9 | 127818309 | A | G | NM_004957 | downstream | dist=4228 | . | . |
| 2049534\|2107421 | 9 | 127818309 | A | G | NR_136302 | ncRNA_intronic | . | . | . |
| 2049534\|2107421 | 9 | 127818309 | A | G | NR_136302 | ncRNA_splicing | c.1378-2A>G | . | . |
| 773598\|704570 | 18 | 31354311 | C | T | NM_001942 | synonymous SNV | c.2115C>T | DSG1 | p.Y705Y |
| 773598\|704570 | 18 | 31354311 | C | T | NR_110788 | ncRNA_intronic | . | . | . |
| 773598\|704570 | 18 | 31354311 | C | T | NR_110788 | ncRNA_splicing | c.298+1G>A | . | . |
| 773598\|704570 | 18 | 31354311 | C | T | NR_110789 | upstream | dist=375 | . | . |

ClinVar
VariationID+AlleleID
As Join Key

Concordance

| key <chr> | NM <chr> | Consequence <chr> | Concordance <chr> | |
|---|---|---|---|---|
| 167569\|178004 | NM_005105 | intronic | intron_variant | |
| 167569\|178004 | NR_002328 | downstream | genic_downstream_transcript_variant | |
| 167569\|178004 | NR_104033 | downstream | genic_downstream_transcript_variant | |
| 167569\|178004 | NR_147182 | ncRNA_intronic | non_coding_transcript_variant | DeDup |
| 167569\|178004 | NR_147182 | ncRNA_splicing | non_coding_transcript_variant | |
| 2049534\|2107421 | NM_000118 | synonymous SNV | synonymous_variant | |
| 2049534\|2107421 | NM_001018078 | downstream | genic_downstream_transcript_variant | |
| 2049534\|2107421 | NM_001114753 | synonymous SNV | synonymous_variant | |
| 2049534\|2107421 | NM_001278138 | synonymous SNV | synonymous_variant | |
| 2049534\|2107421 | NM_001288803 | downstream | genic_downstream_transcript_variant | |
| 2049534\|2107421 | NM_004957 | downstream | genic_downstream_transcript_variant | |
| 2049534\|2107421 | NR_136302 | ncRNA_intronic | non_coding_transcript_variant | DeDup |
| 2049534\|2107421 | NR_136302 | ncRNA_splicing | non_coding_transcript_variant | |
| 773598\|704570 | NM_001942 | synonymous SNV | synonymous_variant | |
| 773598\|704570 | NR_110788 | ncRNA_intronic | non_coding_transcript_variant | DeDup |
| 773598\|704570 | NR_110788 | ncRNA_splicing | non_coding_transcript_variant | |
| 773598\|704570 | NR_110789 | upstream | genic_upstream_transcript_variant | |

Concordance DeDup

| key <chr> | NM <chr> | Consequence <chr> |
|---|---|---|
| 167569\|178004 | NM_005105 | intron_variant |
| 167569\|178004 | NR_002328 | genic_downstream_transcript_variant |
| 167569\|178004 | NR_104033 | genic_downstream_transcript_variant |
| 167569\|178004 | NR_147182 | non_coding_transcript_variant |
| 167569\|178004 | NR_147182 | non_coding_transcript_variant |
| 2049534\|2107421 | NM_000118 | synonymous_variant |
| 2049534\|2107421 | NM_001018078 | genic_downstream_transcript_variant |
| 2049534\|2107421 | NM_001114753 | synonymous_variant |
| 2049534\|2107421 | NM_001278138 | synonymous_variant |
| 2049534\|2107421 | NM_001288803 | genic_downstream_transcript_variant |

### SnpEff annotation output file process

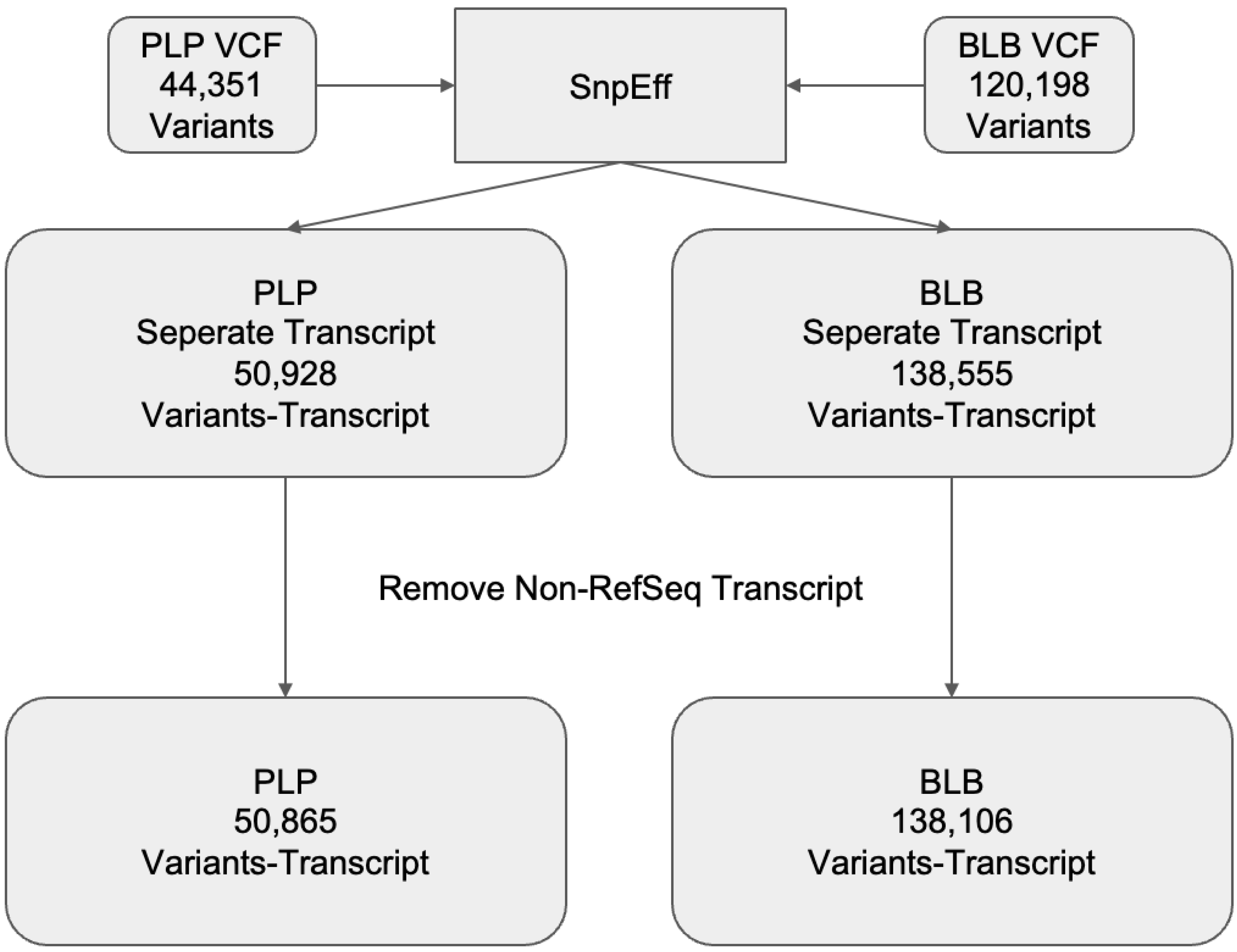


#### SnpEff intergenic_region variant

Note: For intergenic variant, SnpEff does not select the nearest gene as a representative but instead displays gene symbol of both genes. However, because the Feature_ID is not shown as a transcript, it cannot be used for subsequent join comparisons

| Chr | Start | Ref | Alt | Gene_ID | Feature_ID | Feature_Type |
|---|---|---|---|---|---|---|
| 7 | 129774728 | G | A | NRF1-UBE2H | NRF1-UBE2H | intergenic_region |
| 8 | 22161695 | G | C | LGI3-SFTPC | LGI3-SFTPC | intergenic_region |
| 11 | 34916524 | GGGC | G | APIP-PDHX | APIP-PDHX | intergenic_region |
| 11 | 119024925 | G | A | TRAPPC4-HYOU1 | TRAPPC4-HYOU1 | intergenic_region |
| 11 | 119025252 | G | A | TRAPPC4-HYOU1 | TRAPPC4-HYOU1 | intergenic_region |
| 11 | 119026921 | C | T | TRAPPC4-HYOU1 | TRAPPC4-HYOU1 | intergenic_region |
| 19 | 8882863 | C | T | MBD3L1-OR1M1 | MBD3L1-OR1M1 | intergenic_region |
| 22 | 50697269 | T | TG | ARSA-ACR | ARSA-ACR | intergenic_region |
| 22 | 50706113 | C | T | ARSA-ACR | ARSA-ACR | intergenic_region |
| X | 74421488 | G | A | ZCCHC13-SLC16A2 | ZCCHC13-SLC16A2 | intergenic_region |

### VEP annotation output file process

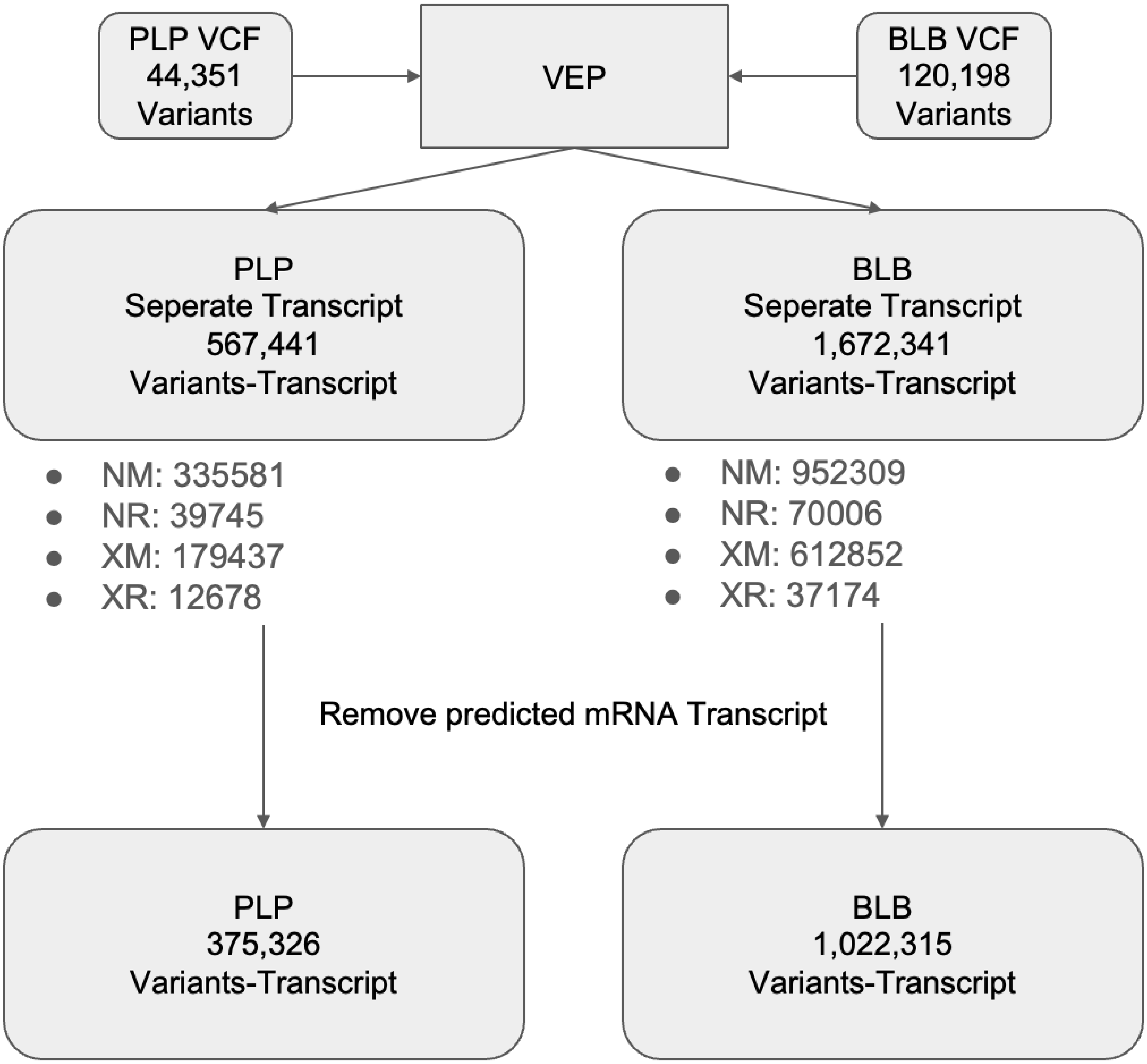
