## Supplementary material for "Toward Automatic Variant Interpretation: Discordant Genetic Interpretation Across Variant Annotations for ClinVar Pathogenic Variants": Key_resource

**Key Resource Table. Software and dataset used in this study.**

| Resource | version | file | reference |
|---|---|---|---|
| ClinVar | January 7, 2024 | VCF, variant_summary_2024-01.txt (GRCh38) | [23] https://ftp.ncbi.nlm.nih.gov/pub/clinvar/vcf_GRCh38/weekly/clinvar_20240107.vcf.gz. |
| bcftools | 1.18 | | [22] |
| Tidyverse | 2.0.0 | | [24] |
| R | 4.3.2 | | [25] |
| ANNOVAR | June 8, 2020 | Transcript set: ensGene(Hg38) 20230315 refGene(Hg38) 20211019 | [14] https://annovar.openbioinformatics.org/en/latest/ |
| SnpEff | 5.2 April 9, 2024 | Transcript set: GRCh38.mane.1.2.ensembl GRCh38.mane.1.2.refseq | [15] https://pcingola.github.io/SnpEff/ |
| VEP | 111.0 November 4, 2024 | Ensembl Release 111 | [16] https://asia.ensembl.org/info/docs/tools/vep/index.html |
| Gendiseak | 2.1.2 | SD2-014A Germline SNV Analysis with New ACMG (GRCh38) | [26] https://www.taigenomics.com/gendiseak/signin |
| VariantValidator | | | [36] https://variantvalidator.org |
